# Neural mechanism underlying task-specific enhancement of motor learning by concurrent transcranial direct current stimulation

**DOI:** 10.1101/2021.01.31.429080

**Authors:** Ying Wang, Jixian Wang, Qing-fang Zhang, Ke-wei Xiao, Liang Wang, Qing-ping Yu, Qing Xie, Mu-ming Poo, Yunqing Wen

## Abstract

The optimal protocol for neuromodulation by transcranial direct current stimulation (tDCS) remains unclear. Using rotarod paradigm, we found that mouse motor learning was enhanced by anodal tDCS (3.2 mA/cm^2^) during but not before or after task performance. Dual-task experiments showed that motor learning enhancement was specific to the task accompanied by concurrent anodal tDCS. Studies using stroke model mice induced by middle cerebral artery occlusion (MCAO) showed that concurrent anodal tDCS restored motor learning capability in a task-specific manner. Transcranial *in vivo* calcium imaging further showed that anodal and cathodal tDCS elevated and suppressed neuronal activity in the primary motor cortex (M1), respectively. Anodal tDCS specifically promoted the activity of task-related M1 neurons during task performance, suggesting that elevated Hebbian synaptic potentiation in task-activated circuits accounts for motor learning enhancement. Thus, application of tDCS concurrent with the targeted behavioral dysfunction could represent a more effective approach for treating brain disorders.

## Introduction

Transcranial direct current stimulation (tDCS) is now widely used for non-invasive modulation of brain functions in healthy subjects and patients with brain disorders, ranging from neurological and psychiatric diseases to stroke-induced dysfunction [1–4]. For example, many previous reports have demonstrated that tDCS applied in the primary motor cortex (M1) can improve motor function of stroke patients [5, 6], but other studies yielded no significant effects [7]. Neuromodulation by tDCS has also been used to alleviate cognitive deficits, such as working memory [8–10], attention [11–13], expression and comprehension of language [14–16], with both positive and negative results. The variability of tDCS effects could be attributed to the large variation in tDCS parameters (current intensity, duration, timing, polarity, stimulation site), electrode configurations, and individual differences among patients. For defining the optimal treatment parameters and protocols, understanding neural mechanisms underlying the tDCS action on the brain is critical. Furthermore, the effects of individual patient’s cranial anatomy on the pattern of current distribution within the brain needs to be considered.

Another important parameter is the timing of tDCS application relative to the patient’s performance of the targeted behavior. In treating motor deficit of stroke patients, anodal [5, 6] or cathodal [5] tDCS was found to produce positive effects on the motor function. Some studies also showed that tDCS combined with the targeted motor task can improve motor function [17, 18]. However, a meta-analysis has shown no conclusive advantage by coupling tDCS with cognitive training as compared to tDCS alone [19]. In this study, we compared specifically the effects of tDCS on mouse motor learning between tDCS that was applied during (“online”) and before or after (“offline”) the motor task training. We found strong evidence that only online anodal tDCS could enhance motor learning, and the effect was task-specific.

Computational modeling studies have predicted the direction and distribution of electric fields in the human brain produced by tDCS, demonstrating that in the human brain the current flows predominantly parallel to the cortical surface [20, 21]. The modeling results also suggest that axon terminals were more susceptible to current-induced polarization than the soma [20]. Measurements of transcranial magnetic stimulation (TMS)-elicited motor evoked potentials indicated that anodal tDCS of human motor cortex for 9-13 min could induce sustained elevation of cortical excitability [22], whereas cathodal tDCS for 9 min caused prolonged inhibition of cortical excitability [23]. Direct current stimulation (DCS) of mouse brain slices has shown that DCS combined with low-frequency synaptic activation (LFS) induced long-lasting synaptic potentiation (LTP), an effect that depended on N-methyl-D-aspartate receptor activation and brain-derived neurotrophic factor [24]. Using *in vivo* two-photon Ca^2+^ imaging to directly monitor cortical activity in the primary visual cortex of urethane-anaesthetized mice, Monai et al [25] found that tDCS activated Ca^2+^ elevation in astrocytes but not in neurons. The mechanism underlying the cell type specificity in the latter study remains unclear. It may be caused by a higher expression of Ca^2+^-sensor in astrocytes [26] or the anaesthetized state of the animal. In the present study, we performed *in vivo* transcranial two-photon Ca^2+^ imaging through the thinned skull to examine neuronal activity in the relatively intact primary motor cortex (M1) of awake mice, particularly the effects of anodal and cathodal tDCS on the activity of M1 neurons related and un-related to the motor task. Our results largely confirmed the excitation and inhibition effects on cortical neurons predicted by computational modeling, and provided a direct mechanistic interpretation of task-specific tDCS effects on motor learning.

The present study examined specifically the notion that modulation of neuronal spiking due to tDCS-induced membrane potential changes [27–29] could be more effective in modulating those neural circuits that are activated at the time of tDCS [2, 29]. Using rotarod running and beam walking paradigms, we examined the enhancement effect and task specificity of online and offline tDCS on motor learning. In both normal wild-type mice and stroke model mice, we found that applying anodal but not cathodal tDCS at the primary motor cortex (M1) during task training markedly enhanced motor learning in a task-specific manner. Together, our findings showed that concurrent application of anodal tDCS with motor task training is more effective in promoting motor learning, and provided mechanistic interpretation of this effect based on cortical neuronal excitation.

## Materials and Methods

### Mice

The primary objective of this study is to examine the neural mechanism underlying tDCS modulation of motor learning. All animal procedures were approved by the Animal Committee of the Institute of the Neuroscience (ION)/Center for Excellence in Brain Science and Intelligence Technology, Chinese Academy of Sciences. For behavioral experiments, male wild-type C57BL/6J mice (7-10 weeks old, from Slyke Co.) were used and randomly assigned to two groups in each experiment: tDCS-treated and sham (0 current)-treated. For MCAO model experiments, male wild-type C57BL/6J mice (8-14 weeks old, male, from Slyke Co.) were used. For *in vivo* two photon imaging of neuronal activity, transgenic mice expressing Thy-1 GCaMP6s (8-14 weeks old, male/female, background strain C57BL/6, purchased from the Jackson Laboratory, Bar Harbor, USA) were used. Mice numbers in each experiment are described in figure legends and main text. Mice were housed under a 12-h light-dark cycle (light during 7 am to 7 pm) at the room temperature (19-22°C) in the ION animal facility. Efforts were made to limit the number of animals used and to minimize their suffering. All behavioral experiments were conducted during daytime at a fixed period during each day for each set of experiments. Two-photon experiments were performed either during daytime at night, due to the availability of the equipment.

### Surgery of electrodes implantation for tDCS

We adopted a unilateral epicranial electrode configuration that was previously used for tDCS in rodents [30]. The stimulation electrode consists of an epicranial implanted tubular plastic jack (inner area 3.14 mm^2^) for behavioral experiments and a circular wire surrounding the chamber above the observation window (area ∼3 mm^2^) for imaging experiments, respectively, with the jack and chamber filled with saline solution (0.9% NaCl) prior to stimulation. The reference electrode was a round tin plate (∼5 mm in diameter) implanted under the contralateral back skin of the neck. For electrode implantation, mice were anesthetized with intraperitoneally (i.p.) injection of pentobarbital sodium (7 mg/kg) and positioned in a stereotaxic frame (model 68030, Reward Co.), the scalp and underlying tissue were removed, and the center of the active electrode was positioned unilaterally on the skull over M1. Stereotaxic coordinates for M1: 0 mm posterior from bregma and 1.5 mm lateral from the midline. During the surgery, the body temperature was maintained at 38°C with a heating pad. All mice were allowed to recover in the cage for 7 days before experiment. tDCS (current: 0.05, 0.1 and 0.2 mA, behavioral experiments; 25 and 50 μA, imaging experiments) was applied to the right M1 with a stimulator (model ST1, Quan Lan Co.). For online tDCS on mice performing beam walking task, costume-made wireless stimulators were used.

### Training for rotarod running task

Mice were familiarized with the experiment room for two hours. A five-lane rotarod (3 cm in diameter, model 47600, Ugo Basile Inc.) was used to assess motor skill acquisition in tDCS-treated and sham-treated mice. Prior to the training period each day, the mouse was given a 5-min familiarization period on the rotarod, with a constant low rotation speed (day 1 & 2, 4 rpm; day 3 & 4, 8 rpm). For each of four consecutive training days, the training was performed at the same time of the day and consisted of three 5-min rotarod running trials (4 to 40 rpm, day 1 & 2; 8-80 rpm, day 3 & 4) [31], interleaved with 5-min rest periods off the rotarod. This procedure is a more sensitive assay for examining motor learning, because the performance of some mice on the easier rotarod (at 4-40 rpm) reached a ceiling at 40 rpm within 2 days, doubling the rotation speed allows mice to show higher extent of motor learning in the following days. In this study, we found that this procedure produced consistent motor learning behavior among different groups of mice and under several different test conditions (e.g., dual motor tasks). Each trial ended when a mouse fell off the rotarod or turn one full revolution, or had reached a duration of 300 s on the rotarod [32]. “Online” tDCS was applied during each trial, and the current stimulation was absent during inter-trial intervals (ITIs). “Offline” tDCS was applied when the animals did not perform the task. Digital video recording was made during the training for later analysis.

### Dual-task training for rotarod running and beam walking

After the training for rotarod running each day (by the same protocol as described above), the mice were allowed to rest for ∼5 h in their home cages before subjected to training for beam walking task. The beam walking training followed that described previously [33], consisting of walking across a 100 cm-long thin beam with a width of 25-, 7-, or 3-mm. The light onset at the start point in the dark room triggered the mouse to walk towards the dark chamber at the other end of the beam. The training was performed over four consecutive days. Each day, a mouse was subjected to a familiarization of 25 mm-wide beam, followed by 3 trials of a beam training (day1&2, 7-mm beam; day 3&4, 3-mm beam). Mice had a 2-min ITI rest in their home cages. A soft cloth was stretched below the beam to protect mice in case of any fall. A video camera was placed on each side of the beam to record the time of crossing and the number of hindlimb slips over a standard 80-cm length on the beam. Slips of both hindlimbs were counted for normal mice, and only slips of the hindlimb contralateral to the lesioned cortex were counted for MCAO model mice.

### Transcranial *in vivo* two-photon imaging

For two-photon imaging, surgery procedure was performed with mice under anesthesia with isoflurane and oxygen mixture, with the body temperature maintained at 38°C with a heating pad. After exposure of the skull, a metal frame was attached to the skull using a dental acrylic, and thinning was performed over a circular region (∼2 mm in diameter) of the skull above the motor cortex (window center site: bregma, 0 mm; mediolateral, 1.5 mm), first by a high-speed micro-drill, followed by thinning of the inner compact bone layer with a microsurgical blade until blood vessels under the skull became clearly visible. Final skull thickness estimated by post-thinning histological measurements was 15.9 ± 0.86 μm (n = 4 mice).

For two-photon imaging, mice were first subjected to 1-day of training on the rotarod, and imaging was then performed on a rotating treadmill (with a constant speed equivalent to the rotarod rotation speed of 15 rpm, 23.6 mm/s), and the animal’s behavior was monitored by an infrared camera. Two-photon imaging was performed with a resonant scanner-based B-Scope (Thorlabs), with the excitation wavelength set at 910 nm (Ti-Sa laser, Spectra Physics) and a field-of-view (FOV) of 350 × 350 μm (512 × 512 pixels) under a 16× objective (NIKON, NA 0.8). Images were acquired using the ThorImage software at a frame rate of 15.6 Hz for 25 minutes or 30 minutes according to different experimental goals. Mice were trained in two behavioral paradigms with tDCS. *First paradigm* (Fig. 3): Mice were running on the treadmill at a constant speed (“task” state) or resting on the treadmill (“rest” state). Measurements of Ca^2+^ signals include 5-min baseline before and after two 5-min tDCS sessions, which were also separated by 5-min baseline (total imaging time 25 min). *Second paradigm* (Fig. 4): For the task state, mice began running on the treadmill following 5-min rest on the treadmill, and 5-min tDCS was applied on M1 after running for 10 min on the treadmill, followed by 10 min running (total running time 25 min, total imaging time 30 min). For the rest state, 5-min tDCS was applied at 5 min after the onset of the experiment on the treadmill, followed by 10 min rest (total imaging time 20 min).

### MCAO

Rodent models of focal cerebral ischemia has been developed to mimic human ischemic stroke, using the procedure of intraluminal suture occlusion of middle cerebral artery (MCA occlusion, MCAO) [34]. This MCAO mouse model has been widely used in studying stroke-induced pathophysiology such as cell death or changes in synaptic structures [35–37] and in designing new prophylactic, neuroprotective, and therapeutic agents [38]. The mice were anesthetized with pentobarbital sodium (7 mg/kg) via i.p. injection, with body temperature maintained at 38°C during surgery. A midline incision was made at the neck and the left common carotid artery (CCA), external carotid artery (ECA), and internal carotid artery (ICA) were identified and ligated. For MCAO, a silicone-coated round-tip MCAO suture (MSMC21B120PK50, RWD Co.) was gently inserted from the ECA stump to the ICA, up to about 10 mm, stopping at the middle cerebral artery (MCA), following the method previously reported [39]. After 90, 60 or 0 min of occlusion, the MCAO suture and ligation were withdrawn. The neck skin was sewn back after blood reperfusion was confirmed.

### TTC (2,3,5-triphenyltetrazolium chloride) staining and Laser Speckle Contrast Imaging (LSCI)

Mice were anesthetized with i.p. injection of pentobarbital sodium (7 mg/kg), and their brains were removed for histology at one day after reperfusion. A series of 2-mm coronal sections were obtained by the brain matrix (model 68707, RWD Co.). The infarct area was shown using 2,3,5-triphenyltetrazolium chloride (2%, Sigma) staining method described previously [40]. In the imaging procedure, the mice were anesthetized with i.p. injection of pentobarbital sodium (7 mg/kg) and a midline incision was made to expose the skull for LSCI before, during and after MCAO, following the previously reported method [41]. The LSCI images before MCAO were used as baseline images. The exposure time for each image was 5 msec and the frame rate was 50.6 frames per second. In the LSCI system (RFLSI III, RWD Co.), the mouse cortex was illuminated by a reshaped laser beam from a 785 nm laser diode. Two hundred speckle images were recorded in each imaging section.

### Quantification in two-photon imaging

For two-photon imaging, the fluorescence signals were quantified by MATLAB-based software (MathWorks) after movement correction of the image stacks with a Turboreg plugin (Image J software, National Institutes of Health) [42]. Fluorescence of the single cell was measured over the region covering each neuronal soma, which was defined by the image stack. Fluorescence change ΔF/F_0_ was defined as (F-F_0_)/F_0_, where F_0_ is the baseline fluorescence averaged over a 5-min period before the onset of the first tDCS. To summarize data from all mice, we obtained the average ΔF/F_0_ during the last 2 min of tDCS by the average values during the 2-min baseline period prior to tDCS for each mouse. For analysis of post-tDCS persistence of activity alteration, we measured the averaged fluorescence changes (ΔF/F_0_) during the last 30 s of every tDCS period and during the subsequent post-tDCS activity at 30-sec bins for 5 minutes.

### Statistics

For behavioral training, rotarod data on “time on rod” and “terminal speed”, and beam walking data on “number of slips” were analyzed by two-way ANOVA. Data for learning rates for rotarod and beam walking were analyzed by two-tailed unpaired *t*-test. For two-photon imaging data, significance tests were performed between data obtained during anodal/cathodal tDCS and baseline (2 min before each tDCS onset) using two-tailed paired t-test. The statistical analysis was performed using commercial software (GraphPad Prism, Version 5.0, GraphPad, San Diego, USA). Data were considered significant as follows:“*”, p< 0.05; “**”, p< 0.01.

## Results

### Online anodal tDCS enhances mouse learning of rotarod running task

Mice were subjected to a rotarod running task that began each day with a 5-min familiarization period for rotarod running at a constant low rotation speed, followed by three 5-min trials with gradually increasing speed (4 to 40 rpm, day 1 & 2; 8-80 rpm, day 3 & 4) [31] that were spaced with 5-min ITIs off the rotarod (Fig. 1A). Mice were subjected to tDCS at designated time with anodal (“+”) or cathodal (“-”) currents, or without current (sham control “S”) (Fig. 1B). The mouse normally learned well in running on the rotarod over four training days, as shown by the increasing duration of staying on the rotarod (Fig. 1C) and increasing terminal rotor speed when the mouse dropped off the rotarod (Fig. 1D). When tDCS was applied to the right primary motor cortex (M1) during the familiarization period and all three task trials each day (“online” stimulation), we found a significant increase in both the time on the rotarod and the terminal speed, beginning on the second day of training (Fig. 1C, Online, n=13 mice; Sham, n=10 mice, and movie S1, S2). This enhancement of motor learning remained detectable at the 14 but not 28 day after training (fig. S1, A, B; same n as above). The results were further quantified by the rate of learning, as defined by the normalized difference of the terminal speed between the first and last training trials (Fig. 1E; same n as above). Doubling the anodal tDCS current magnitude to 0.2 mA caused occasional convulsion in mice, and reducing current magnitude to 0.05 mA resulted in no learning enhancement (Fig. 1E, and fig. S2A, B; n=11 for both Online and Sham). We thus chose 0.1 mA for the standard anodal tDCS in this study. Furthermore, we found no enhancement of rotarod learning when the same online anodal tDCS was applied to the primary visual cortex (V1, Fig. 1E, and fig. S3A, B; Online, n=11; Sham, n=12), indicating stimulation site-specific tDCS effect. The rotarod learning was not affected by the procedure of surgery and electrode installation, as shown by comparing the same motor learning of mice that were not subjected to the procedure (fig. S4A, B; Surgery, n=9; Control, n=12).

**Fig. 1.**
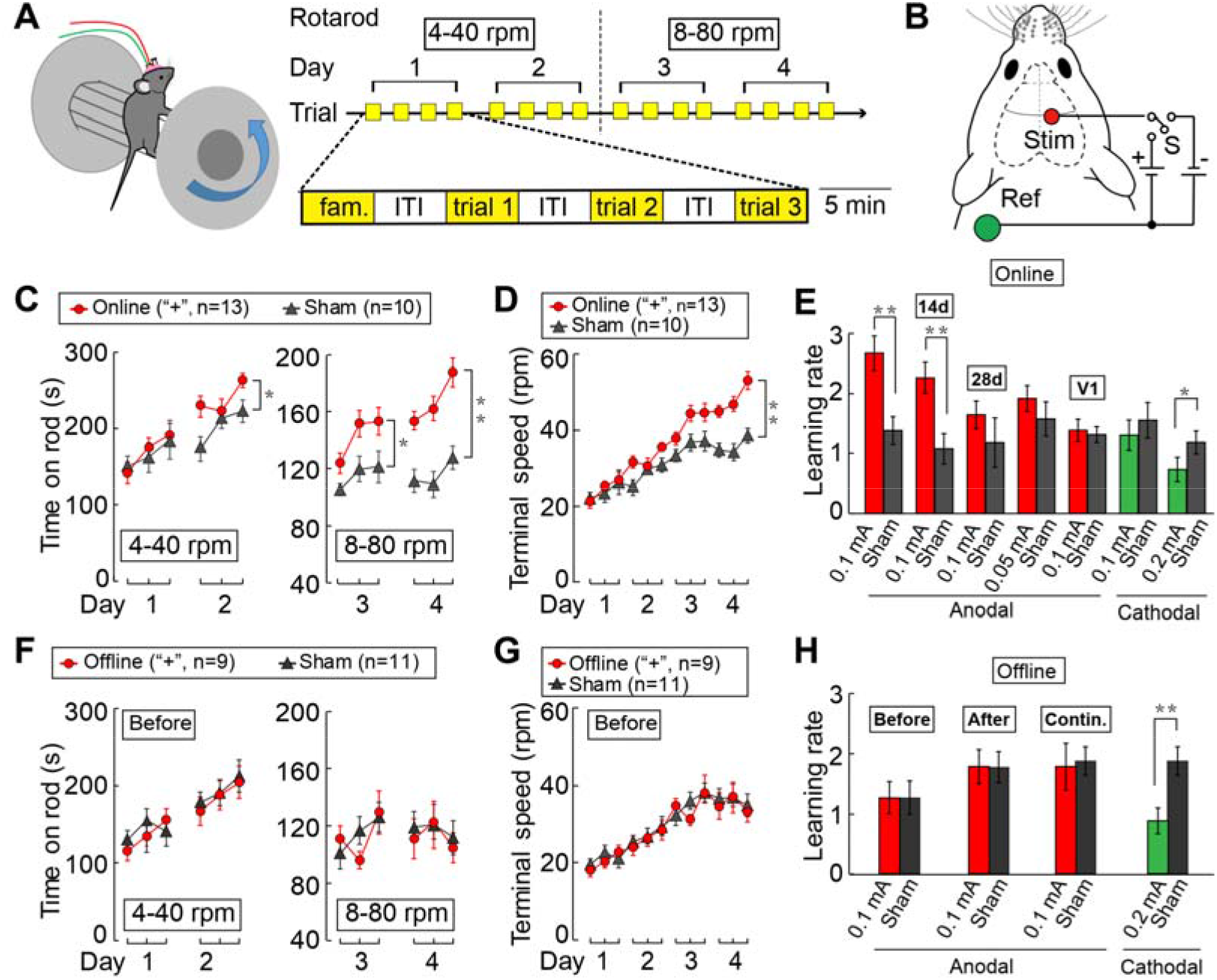
Effects of tDCS on mouse learning of the rotarod running task. (**A**) Training Protocol: The mouse was subjected each day to a 5-min familiarization trial at a constant low speed, followed by three 5-min trials (separated by 5 min inter-trial interval) at linearly increasing rotation speed. (day 1 & 2: 4 - 40 rpm; day 3 & 4: 8-80 rpm). (**B**) Schematic diagram depicting the electrode configuration. “**Stim**”: tDCS electrode. “**Ref**”: reference electrode. “**S**”: sham (no current). “**+**”: anodal. “**-**”: cathodal. (**C**) The average time of staying on the rotarod during each trial. (**D**) The terminal rotation speed at which mice fell off the rotarod during each trial. “**Online**”: anodal tDCS (0.1 mA) was applied during each trial. “**n**”: total number of mice examined. (**E**) Summary of results showing the learning rate, as defined by the normalized difference of terminal speed between the last and the first trial of the entire training period. Data depict standard 4-day training with (color bars) and without (sham, black bars) online anodal (or cathodal) tDCS, applied to M1 at different current amplitudes. “**14d**” and “**28d**” refer to results obtained with 3 additional training trials at 14 and 28 d after training. “**V1**”: the tDCS was applied to V1 instead of M1. (**F**-**H)** The results from experiments in which tDCS was applied during ITIs, presented in the same manner as those described above for C-E. “**Before**” and “**After**” refer to the average values obtained with tDCS applied during ITIs before and after each trial, respectively. “**Contin.**”, 20-min continuous tDCS applied before the familiarization trial. Error bars, SEM. Significant difference was found between the data sets connected by lines (“*”, p< 0.05; “**”, p< 0.01; C, D, F, G: two-way ANOVA; E, H: unpaired *t* test).

In contrast to the learning enhancement described above, we found that anodal tDCS (at 0.1 mA) applied during all 5-min ITIs before or after rotarod running (“offline” stimulation) had no effect on the rate of rotarod learning (Fig. 1F-H, and fig. S5A, B; “After”: Offline, n=12, Sham: n=11). Furthermore, no effect was found when anodal tDCS was applied continuously for 20 min before the task onset (Fig. 1H, and fig. S5C, D; “Contin.”: Offline: n=12, Sham: n=11), a protocol often used in clinical research [43]. In contrast to anodal tDCS, online *cathodal* tDCS (0.1 mA) at M1 also had no effect on rotarod learning (Fig. 1E, and fig. S6A, B; Online: n=7, Sham: n=5). However, when the cathodal current was increased to 0.2 mA, learning was impaired at the 3^rd^ and 4^th^ day of training (Fig. 1E, and fig. S6C, D; Online: n=8, Online sham: n=8). Unlike that found for anodal tDCS, both online and offline cathodal stimulation at 0.2 mA resulted in similar impairment of learning (Fig. 1E, H, and fig. S6C, D). As shown later, this may be attributed to the long-lasting (>5 min) suppression of neuronal firing by cathodal tDCS. Taken together, these findings show that tDCS could bi-directionally modulate rotarod learning, and that the enhancement effect was significant only when concurrent anodal tDCS was applied with the performance of the rotarod task.

### Task-specific enhancement of motor learning by anodal tDCS

The effect of online anodal tDCS on motor learning may be attributed to specific enhancement of rotarod running skill or improvement of motor coordination in general. To address this issue, we introduced a beam-walking learning task, in which the mouse was given a short familiarization period for walking along a wide beam (25 mm in width), followed by 3 trials of walking on a narrow beam each day (7 mm, day 1 & 2; 3 mm, day 3 & 4) (Fig. 2A). The learning process was shown by a gradual reduction of the mean number of hindlimb slips and the mean transverse time during beam walking, and the learning rate was quantified by the normalized difference of the mean number of slips between the last and the first beam-walking trial on the 3-mm beam over the 4-day training period.

**Fig. 2.**
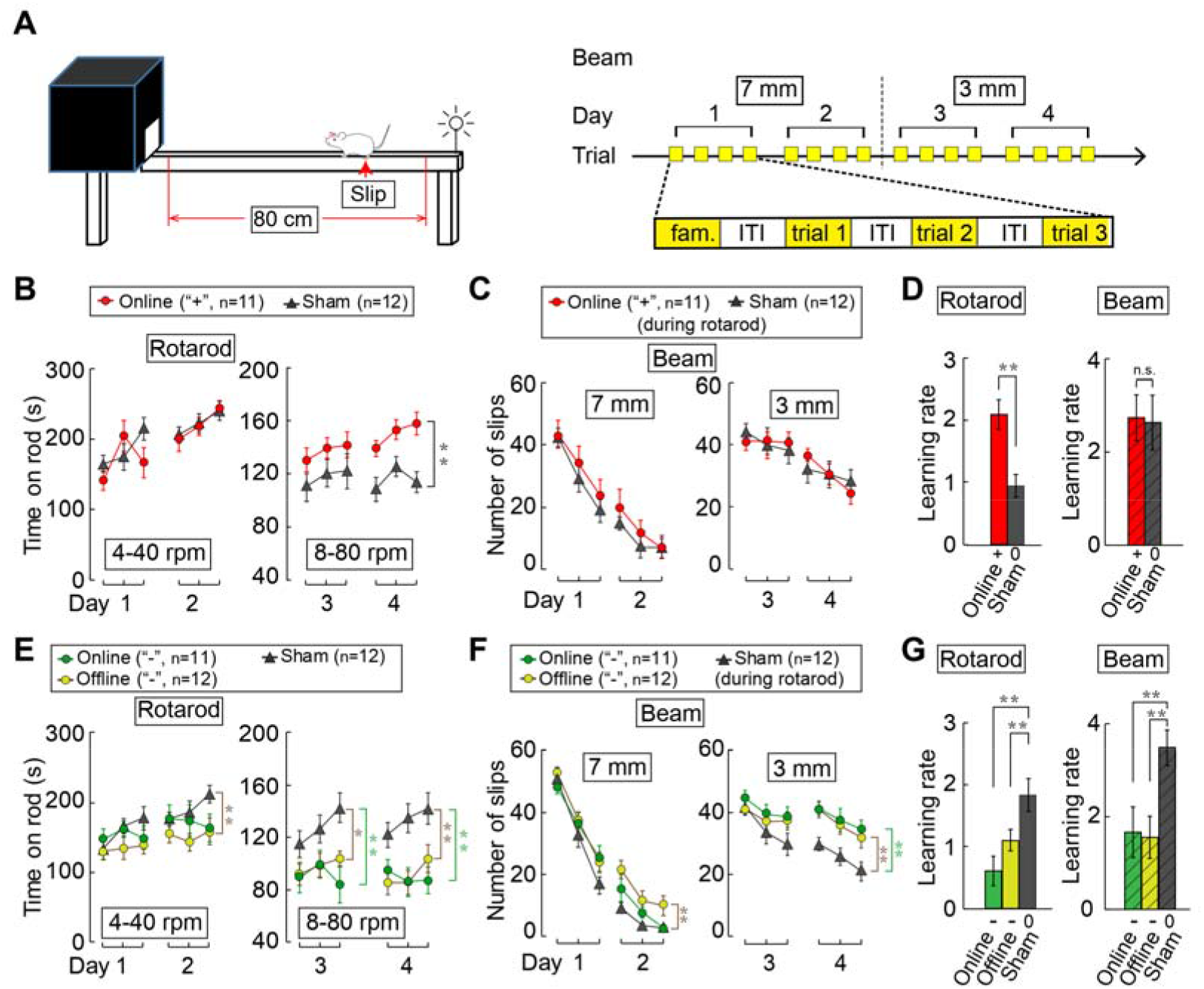
Effects of tDCS-induced modulation of rotarod learning on the learning of beam walking. (**A**) Experimental protocol of beam walking. The mice were subjected to anodal online tDCS in the same manner as that described in Fig. 1C, except that rotarod task was followed by a beam walking learning task in the absence of tDCS. The mice were subjected to wide beam (25 mm) familiarization, followed by three trials on the thinner beam (day 1 & day 2, 7 mm; day 3 & day 4, 3 mm). (**B**, **C**) Data from dual-task experiments. Average time on the rotarod (in B) was presented as that in Fig.1C. The reduction in the average frequency of hindlimb slips (in C) during the 4-day training of beam walking. Note that online anodal tDCS during rotarod running improved rotarod learning (in B), but had no effect on learning beam walking (in C). “**n**”: total number of mice examined. (**D**) Summary of results showing rotarod learning rate and beam walking learning rate, as defined by normalized difference of the slip frequencies between the last and the first trial of the beam walking on the 3-mm beam. (**E-G**) Results of rotarod learning and beam walking learning by cathodal online (or offline) tDCS during rotarod learning. “+”: anodal tDCS; “-”: cathodal tDCS; “0”: no current. Error bars, SEM. Significant difference was found between the data sets connected by lines (“*”, p< 0.05; “**”, p< 0.01; B, C, E, F: two-way ANOVA; D, G: unpaired *t* test).

In the first set of experiments, we measured beam-walking ability before and after 4 d of rotarod training, and the beam walking ability was not affected by rotarod training, as reflected by similar reduction of hindlimb slips as that found in untrained mice (fig. S7A-C; Rotarod: n=10, Control: n=12). This implies that motor learning was specific to the trained motor task. In the second set of experiments, we trained the mice to perform both rotarod running and beam walking (“dual tasks”) each day over four training days, and found that rotarod learning did not affect the learning rate for beam walking, which was comparable to that resulted from beam-walking training alone (fig. S8A-C; Rotarod: n=10, Control: n=12). Thus, there was no transfer of learning from rotarod running to beam walking. Importantly, when we enhanced the rotarod learning by the online anodal tDCS, the learning rate for beam walking was not affected in the dual-task training (Fig. 2B-D; Online: n=11, Sham: n=12). Conversely, when the learning of beam walking was enhanced by online anodal tDCS (fig. S9A, B, and movie S3-6; Online: n=15, Sham: n=15), we found no enhancement of learning for rotarod running (fig. S10A-D; Online: n=18, Sham: n=17). Thus, online anodal tDCS during a specific task did not lead to general enhancement of motor learning. In contrast to this specific anodal tDCS effect, we found that both online and offline *cathodal* tDCS during rotarod training had suppressive effects on learning both rotarod running (Fig. 2E, G; Online: n=11, Offline: n=12, Sham: n=12) and beam walking (Fig. 2F, G; Online: n=11, Offline: n=12, Sham: n=12).

### Modulation of neuronal activity by anodal and cathodal tDCS

We next examined the action of tDCS on the activity of M1 neurons using transcranial *in vivo* two-photon calcium imaging. We used *thy-1* transgenic mice expressing Ca^2+^-sensitive fluorescent protein GCaMP6s in cortical neurons, and monitored spiking activity of individual neurons by measuring the elevation of GCaMP6s fluorescence [44] through the skull after a skull thinning procedure (see Methods). The activity of cortical neuron populations in the layer 2/3 of M1 was recorded in head-fixed mice on a treadmill that alternated between “task” (during mouse running on the steadily moving treadmill, at velocity 23.6 mm/s) and “rest” (during mouse resting on the stationary treadmill, at zero velocity) states (Fig. 3A). We observed substantial spontaneous activity of M1 neurons, as reflected by pulsatile changes of fluorescence signals (Fig. 3B, movie S7), which are known to correlate with spiking rate of the neurons [44, 45]. When anodal tDCS was applied through a saline pool above the thinned skull, we observed a gradual increase of fluorescence signals in many neurons (movie S7). Figure 3C (n=6 cells) illustrates changes of fluorescence signals (ΔF/F_0_) in 6 example neurons (boxed in Fig. 3B) during the task and rest periods when two consecutive anodal or cathodal tDCS were applied (each for 5 min). Apparent elevation of Ca^2+^ activity by anodal tDCS (25 µA) was observed in 4/6 neurons during the task but not the rest period, and all 6 neurons showed strong inhibition of the activity during cathodal tDCS (50 µA) (Fig. 3C). The same group of cells were monitored before and after two episodes of anodal and cathodal tDCS sequentially, under the task and rest conditions.

The reproducibility of tDCS effects on neuronal activity was examined in separate experiments on eight mice where either anodal or cathodal tDCS was repeated after an interval of 5 minutes. The results are summarized in Fig. 3D for all cells in the imaged field. Significant elevation and suppression of fluorescence signals were induced by anodal and cathodal tDCS during the task period, respectively (Fig. 3E, F; n=8 mice). We also noted that changes in the average fluorescence signal subsided gradually after each tDCS offset, and that the suppressive effect of cathodal tDCS persisted for a longer duration than the enhancement effect of anodal tDCS (Fig. 3G; n=8 mice). This may account for the offline suppressive effect on the rotarod learning described above using only 5-min ITI in the present paradigm.

The M1 neurons monitored in the above experiments may include neurons that were activated for performing the treadmill running task and those unrelated to the task. We thus further inquired whether the tDCS effects differ between these two types of neurons. The activity of all GCaMP6s-expressing M1 cells within the field of view were monitored for 5 min before the task onset to obtain the baseline activity (Fig. 4A). Task-related and un-related cells were defined by their peak fluorescent signal (ΔF/F_0_) within the first 2-min window after the task onset that was above the level of baseline + 1.5 SD and below the level of baseline + 0.5 SD, respectively. Data of all task-related cells (“+”, n=247 cells; “-”, n=158 cells) and task-unrelated cells (“+”, n=22 cells; “-”, n=54 cells) identified in 4 mice were summarized by the activity heatmap and average activity profiles (Fig. 4A). We found during the task period, anodal tDCS induced highly significant elevation of activity in task-related cells, but not in task-unrelated cells. By contrast, the same anodal tDCS of this population of neurons during the rest period had no significant effect on either type of cells (Fig. 4A, B). The inhibitory effect of cathodal tDCS, however, was highly pronounced during both task and rest periods in all neurons (Fig. 4A, B). These results support the notion that the specific effect of anodal tDCS on motor learning was due to the elevation of the activity of task-related neuron circuits.

Taken together, these results support the notion that anodal and cathodal tDCS modulate neuronal firing by inducing depolarization and hyperpolarization of cortical neurons, respectively, consistent with previous findings on isolated brain slices [24, 46, 47]. When applied at the time of specific motor circuit activation, as that occurred during motor task, anodal tDCS could facilitate learning-associated modification of specific motor circuits in M1, via enhancing correlated firing that induces Hebbian long-term potentiation (LTP) of synapses within these circuits.

### Task-specific restoration of motor learning in stroke mice by tDCS

Meta-analyses have shown high variability in the clinical efficacy of tDCS in treating stroke patients [48, 49]. This variability could be attributed in part to differences in the tDCS protocol and individual stroke conditions. In this study, we have examined the effect of tDCS on motor learning in a relatively defined mouse model of stroke. A standard middle cerebral artery occlusion (MCAO) for 60 or 90 min in the mouse’ left hemisphere induced a large lesion within the left somatosensory cortex and part of the motor cortex at one day after MCAO (Fig. 5A). When these mice were subjected to rotarod learning at 14 days after MCAO (Fig. 5A), we found their motor coordination was significantly impaired, as shown by an overall reduction in the time on the rotarod and the rate of rotarod learning, as compared to control mice that underwent MCAO surgery without sustained artery occlusion (Fig. 5B, C; MCAO: n=11 mice, Control: n=12 mice). Furthermore, online anodal tDCS at the left perilesional M1 region (Fig. 5A) largely restored the mouse’ learning of motor coordination and rotarod running (Fig. 5B, C, E; and movie S9, 10; MCAO: n=11, MCAO/Online: n=11). In contrast, offline *anodal* tDCS (Fig. 5F and fig. S11A, B; MCAO/Offline, n=11, MCAO, n=11), online *cathodal* tDCS (Fig. S12A; MCAO/Online, n=8; MCAO, n=9), and offline *cathodal* tDCS (Fig. S12B; MCAO/Offline, n=7; MCAO, n=9) at the same site all had no effect on learning motor coordination and rotarod running in MCAO mice.

**Fig. 3.**
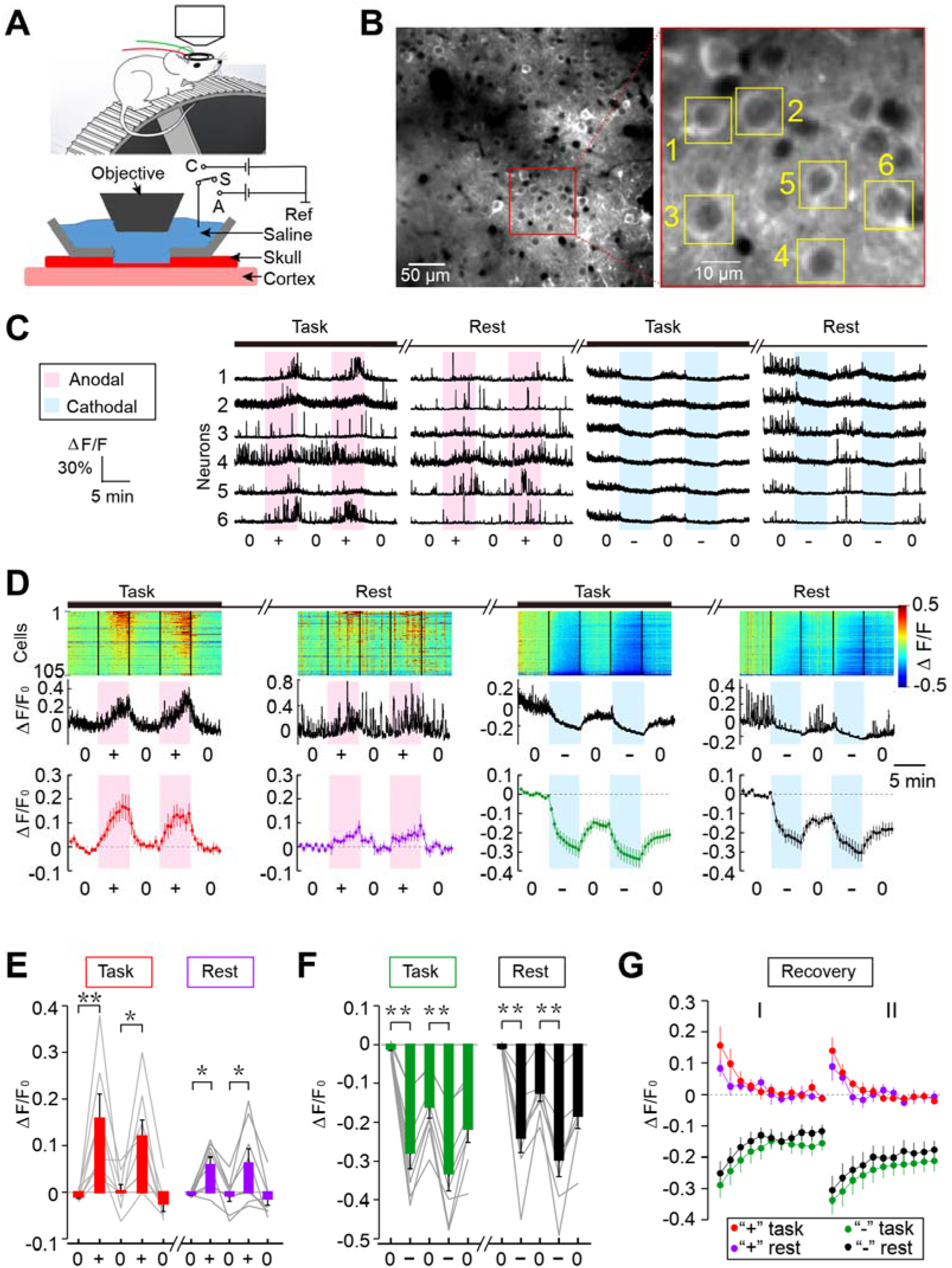
Transcranial two-photon imaging of tDCS-induced modulation of cortical neuronal activity. (**A**) Schematic diagram depicting the optical window over the thinned skull, for two-photon imaging of M1 neurons in a head-fixed mouse on the treadmill that moved at a constant speed during the task. (**B**) Example images of Thy1-GCaMP6s-expressing neurons in M1, viewed through the imaging window. Red-boxed region in the left image is shown at a higher resolution, revealing GCaMP6s fluorescence of individual layer 2/3 neurons. (**C**) Changes of GCaMP6s fluorescence (ΔF/F_0_) with time monitored at six M1 neurons (marked by yellow boxes in B). Pink and blue shading mark the duration of anodal tDCS (at 25 µA) and cathodal tDCS (at 50 μA), respectively. (**D**) Fluorescence changes of all fluorescently labelled cells within the imaged field, recorded from one example mice. The amplitude of ΔF/F_0_ for each cell with time was color-coded with the scale shown on the right. The cells were ordered according to the peak values of ΔF/F_0_. Two panels of traces below represent average ΔF/F_0_ for all cells shown above, and the average ΔF/F_0_ for all cells from 8 mice, respectively. (**E**, **F**) Summary on tDCS-induced GCaMP6s fluorescence changes for data from all mice (n=8). Average fluorescence changes (ΔF/F_0_) during the last 2-min of tDCS were normalized by the average values during the 2-min baseline period prior to tDCS, for two consecutive trials under the task and rest conditions. Data for the same set of neurons in each mouse were connected by lines. Significant difference was marked (“*”, p < 0.05; “**”, p < 0.01; paired *t* test). (**G**) Post-treatment persistence of tDCS effects was shown by the average fluorescence changes with time, normalized by the values at the time of termination of anodal or cathodal tDCS, for task and rest conditions. Error bars, SEM.

**Fig. 4.**
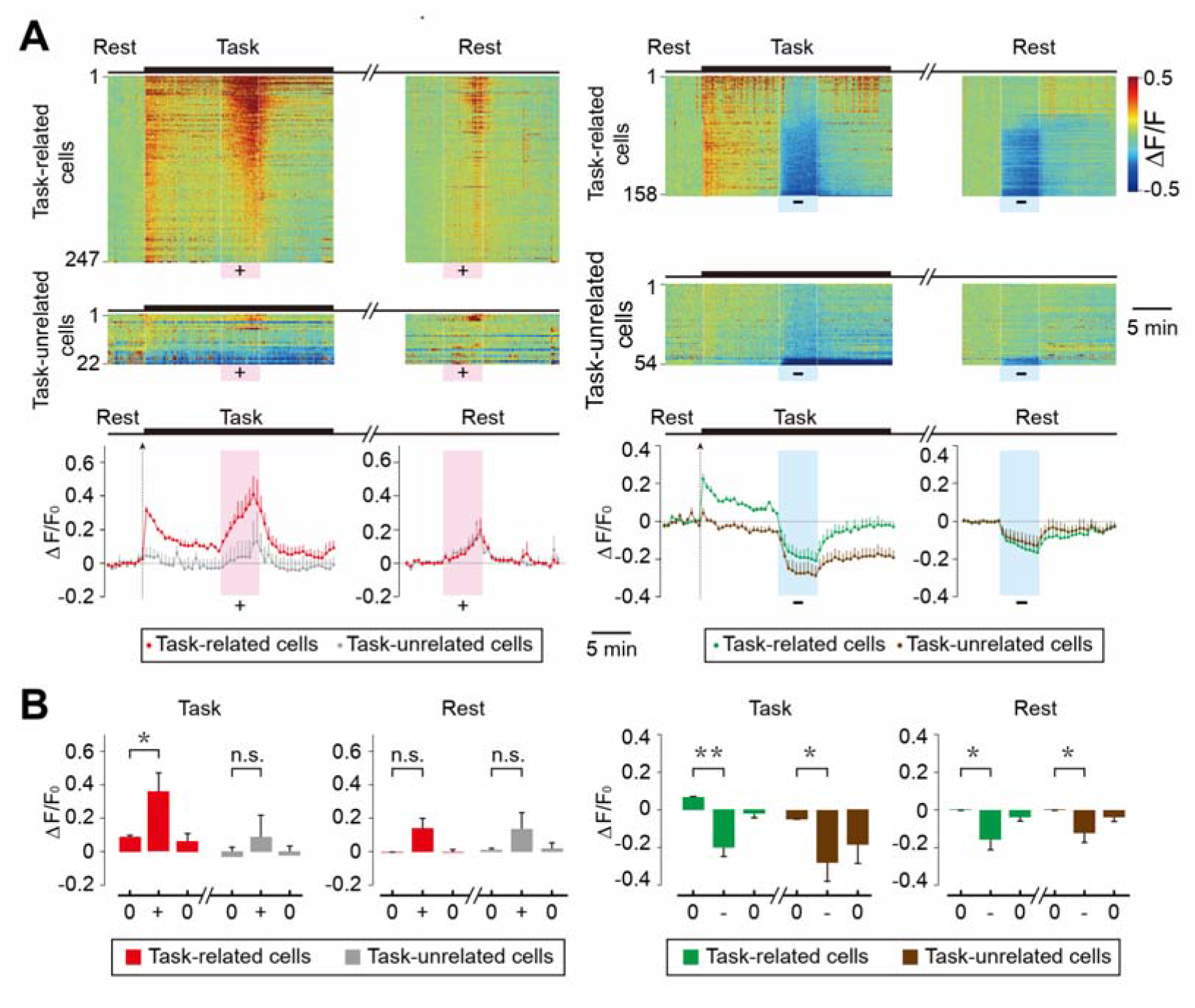
Modulation of activity of task-related and task-unrelated cortical cells by tDCS. (**A**) Fluorescence changes (ΔF/F_0_) of task-related cells and task-unrelated cells within the imaged field (definitions in Methods) were shown by activity heat maps of M1 cell populations. The amplitude of ΔF/F_0_ was normalized for each cell by the baseline during 5-min period before the task onset and color-coded with the scale shown on the right. All cells (anodal: n=269; cathodal: n=212) recorded from 4 mice were grouped and ordered according to the peak values of ΔF/F_0_ within the tDCS time window. Curves below depict changes in the average ΔF/F_0_ with time during the experiment shown above, for task-related and task-unrelated cells. Error bars, SEM. (**B**) Summary of tDCS-induced ΔF/F_0_ for data from all 4 mice. Average ΔF/F_0_ during the tDCS period (“+” or “-”) were compared with those during the period before and after tDCS (“0”). Average ΔF/F_0_ during the last 2 min of each period were used for the histogram. Error bars, SEM. Significant differences are marked by “*”(p < 0.05) or “**”(p < 0.01), and no significance marked by “n.s.” (paired *t* test).

**Fig. 5.**
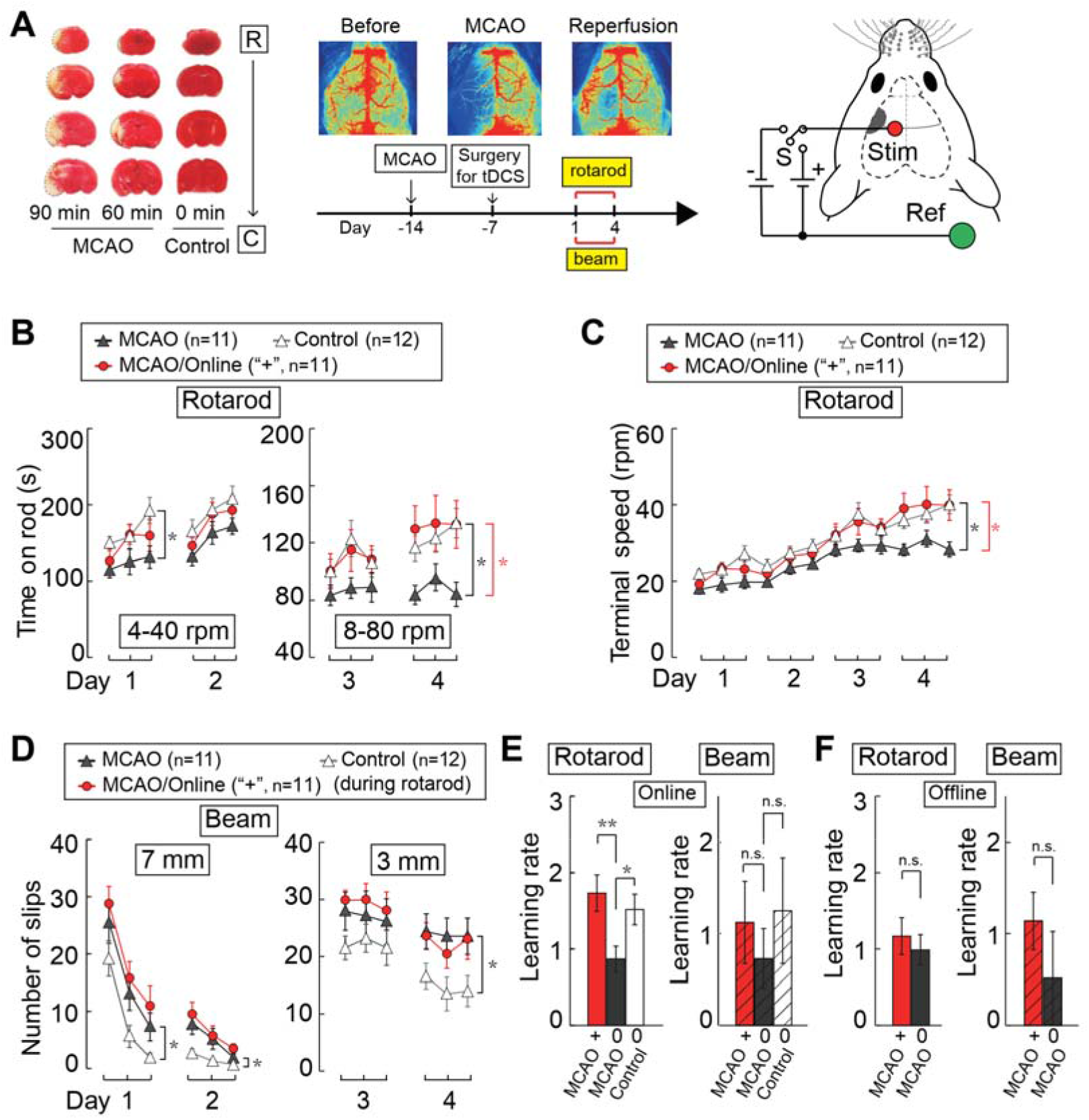
Task-specific restoration of motor learning ability by online anodal tDCS in MCAO mice. (**A**) TTC staining (left) and laser speckle contrast imaging (LSCI) (right) showing the lesion induced by MCAO, which was performed for 90, 60 or 0 min prior to reperfusion. The time schedule of MCAO, surgery for tDCS, and training for rotarod running and beam walking is shown. Schematic diagram on the right depicts the placement of tDCS electrodes in MCAO mice. Infarct area marked in gray, and the stimulation electrode (“**Stim**”) covered parts of M1 and somatosensory cortex. (**B**, **C**) The average time on and terminal speed of the rotarod, for MCAO mice treated with online anodal tDCS, sham-stimulation, and sham-MCAO surgery (control) in dual-task experiments, in which tDCS was applied only during rotarod running. Data presented in the same manner as in Fig.1, C and D. “**MCAO**”: mice subjected to 90-min occlusion in MCA; “**Control**” (Sham-MCAO): mice subjected to the same surgery with no occlusion in MCA; “**Online**”: MCAO mice subjected online tDCS during rotarod running. “**n**”: total number of mice examined. (**D**) The average frequency of hindlimb slips (contralateral to the lesion) during beam walking. (**E**) Learning rate for rotarod running and beam walking in MCAO mice subjected to online anodal tDCS. (**F**) Learning rate of rotarod and beam walking in MCAO mice that were subjected to offline anodal tDCS. “**Offline**”: MCAO mice subjected tDCS before rotarod running. “**+**”: anodal tDCS; “0”: no current. Error bars, SEM. “n.s.”, no significance. Significant difference was found between the data sets connected by lines (“*”, p< 0.05; “**”, p< 0.01; B, C, D: two-way ANOVA; E, F: unpaired *t* test).

In the absence of tDCS, 90-min MCAO impaired motor learning of both rotarod running and beam walking, as compared to control mice (Fig. 5B-E; MCAO: n=11; Control, n=12). However, the mice that showed rotarod learning restoration by online anodal tDCS did not improve the learning of beam walking, as compared to those subjected to sham tDCS treatment during rotarod running (Fig. 5B-E; MCAO: n=11, MCAO/Online: n=11). In contrast, offline anodal tDCS during rotarod training had no effect on learning both rotarod running and beam walking (Fig. 5F and fig. S13A-D; MCAO: n=12, MCAO/Offline: n=14). Therefore, the restoration of rotarod learning in MCAO mice by anodal tDCS was task-specific, rather than a general restoration of motor learning. Based on the above finding of elevated neuronal firing induced by anodal tDCS, the restoration of rotarod learning may involve specific enhancement of residual neural circuits after MCAO that were activated during rotarod running, without affecting those underlying beam walking.

## Discussion

The timing of tDCS relative to the targeted task performance has been addressed in previous studies of healthy human subjects and stroke patients, but conflicting results have been reported, as summarized by various meta-analyses [48, 49]. For examples, online but not offline anodal tDCS of M1 during motor sequence task was found to enhance motor learning, while online cathodal tDCS had no or opposite effects [50, 51]. However, another study using offline anodal tDCS prior to the motor task in human subjects have shown an enhancement effect on motor learning [52]. In cases of prolonged tDCS, the effects on human motor cortex could last for hours [22] and even days [53], and the timing of tDCS becomes less relevant. A previous study using mouse brain slices show that only DCS coupled with low-frequency synaptic activation could induce a long-lasting synaptic potentiation [24]. Direct current stimulation time-locked to the expected onset of low-frequency oscillations (LFO; <4 Hz) could also significantly improve skilled reaching in stroke model rats [54]. Our present results further underscore the importance of concurrent application of neuromodulation during task performance, especially when brief episodes of stimulation was used.

Previous studies on healthy human subjects have shown that anodal tDCS enhanced cognition or motor learning [55–58] and these effects were specific to different level of task difficulty [59, 60] or the site of tDCS [58, 61]. We found that anodal tDCS on M1 specifically enhanced the learning of rotarod task, without affecting the learning of beam walking. Thus, even within the motor domain, concurrent tDCS could exert modulation of specific motor functions. The mechanism underlying the task-specific tDCS effect was further examined in the present study using *in vivo* imaging of M1 neuronal activity. We showed that task-related M1 neurons are preferentially elevated by anodal tDCS, as compared to task-unrelated neurons, during the performance of the motor task. Thus, task-related circuit activation and potentiation account for the increase of motor functions induced by anodal tDCS. The same mechanism also accounts for the effect of low-frequency epidural alternating current stimulation (ACS) in improving grasping dexterity in macaque monkeys after lesion-induced stroke, where ACS was shown to increase co-firing within task-related neural ensembles in the perilesional cortex [62]. Similarly, in chronic stroke patients, tDCS combined with locomotor training with a robotic gait orthosis improved motor restoration [63].

The tDCS current density (3.2 mA/cm^2^) used in the present study was smaller than that used (5.7 mA/cm^2^) by Pedron et al. [30] for studying rat addictive behavior and working memory. This current density is 3-4 times lower than the upper limit of safe tDCS current determined in a rat study [64]. Cathodal tDCS at the 5.7 mA/cm^2^ was also found to improve working memory and skill learning in rats [65]. Similar tDCS current levels were also used in rats for treating status epilepticus (5.7 mA/cm^2^) [66], for promoting recovery from stoke-induced cognitive impairments (2.8 mA/cm^2^) [67], and for elevating dopamine release in the striatum (3.2 mA/cm^2^) [68]. In the previous *in vivo* Ca^2+^ imaging study on astrocyte activation by tDCS [25], the current density was 5.0 mA/cm^2^, similar to the level used in our study. Notably, the standard current density applied to humans (0.029 and 0.057 mA/cm^2^) [43, 69] is much lower than that used in rodent studies. The difference may be attributed to safety consideration, the effectiveness of current penetration through the skull and cortex, electrode configuration, the extent of neuronal activity induced by the current, and the complexity of neural network.

The exact current density induced by tDCS in the cortex remains unclear. In our behavioral study, the effective current density of anodal tDCS was 3.2 mA/cm^2^ at the surface of the intact skull. Histological measurements of the thickness of thinned skull for mice used in our Ca^2+^ imaging experiments yielded an average thickness of 15.9 ± 0.86 μm (n=4 mice). Thus, the average current density was estimated to be ∼0.8 mA/cm^2^ at the observation window (∼2 mm in diameter) for the anodal current applied (25 µA), with a higher density near the center due to non-uniform current distribution. More precise estimate of the effective current density requires further analysis of the pattern of subdural currents, which depend on the electrode configuration and the resistance of various tissues.

We found that application of anodal tDCS to mouse M1 elevated cortical neuronal activity whereas cathodal tDCS suppressed their activity. This mechanism could underlie the tDCS effects on human motor cortex, where anodal and cathodal tDCS increased and reduced corticospinal excitability (revealed by TMS-induced MEP amplitudes), respectively [22, 23, 70]. However, another study using cathodal tDCS of the human motor cortex showed a significant increase and decrease of corticospinal excitability at the total current level of 2 mA and 1 mA, respectively [71]. While the cause remains unclear, this finding underscores the importance of precise control of the magnitude of tDCS current. The tDCS acts by altering neuronal membrane potentials, and currents of different levels could activate or inhibit distinct populations of neurons that have differential firing thresholds, leading to disparate functional effects.

In this study, task specificity was found in the enhancement effect of anodal tDCS on motor learning, but not in the suppression effect of cathodal tDCS. This difference may result from our specific experimental paradigm, in which we used 5-min inter-trial interval between sequential cathodal tDCS. Imaging experiments showed that this short interval did not allow complete recovery of neuronal activity following cathodal tDCS, thus producing offline inhibition effect. By further adjustment of the inter-trial interval, it is possible that task-specific suppression effect could also be achieved by cathodal tDCS.

## Conclusions

In this study, we have characterized the mechanism of action and the appropriate paradigm for the use of anodal tDCS in enhancing motor learning in normal mice and stroke model mice. Our results suggest that concurrent application of tDCS with the performance of the targeted task could elevate the therapeutic efficacy. Our imaging results provide the neuronal mechanism underlying the effect of concurrent tDCS in promoting task performance. This approach of concurrent neuromodulation could be applied to the treatment of other brain disorders, such as obsessive compulsive disorder, auditory hallucination in schizophrenia, epilepsy, and addiction. While the exact neural circuit abnormality of many brain disorders remains to be identified, neuromodulation applied during voluntary or triggered disorder-associated behaviors could help to potentiate or suppress the underlying neural circuits, leading to therapeutic effects.

## Acknowledgements

We thank Drs. Yang Dan, Chun Xu, Liping Wang, Huatai Xu, Zhiqi Xiong for suggestions; Dr. Huanhuan Zeng for technical support; Dr. Zhijie Wang for providing the head-fixation holder and plates. This work was supported by the Strategic Priority Research Program of the Chinese Academy of Sciences (XDB32070100); Shanghai Municipal Science and Technology Major Project (2018SHZDZX05); the Shanghai Key Basic Research Project (18JC1410100); Lingang Lab (LG202106-04-03, LG202105-01-07); Shanghai Pilot Program for Basic Research – Chinese Academy of Science, Shanghai Branch (JCYJ-SHFY-2022-010).

## Author Contributions

Y.W., M.P, and Y.W. designed the study, Y.W., J.W. and L.W. performed the experiments; Q.Z., K.X., Y.W. and Y.W. wrote the data processing program and provided technical help; Q.X. assisted with the design of experiment; Q.Y. helped managing the mouse colony; and Y.W, M.P., and Y.W. wrote the manuscript.

