## Supplementary information for "Neural mechanism underlying task-specific enhancement of motor learning by concurrent transcranial direct current stimulation"

**This supplementary information file includes:**

Figs. S1 to S14

Captions for movies S1 to S10

Two-Sample T-Tests Allowing Unequal Variance

**Other Supplementary Materials for this manuscript include the following:**

Movies S1 to S10

**Supplementary figures (1-14)**

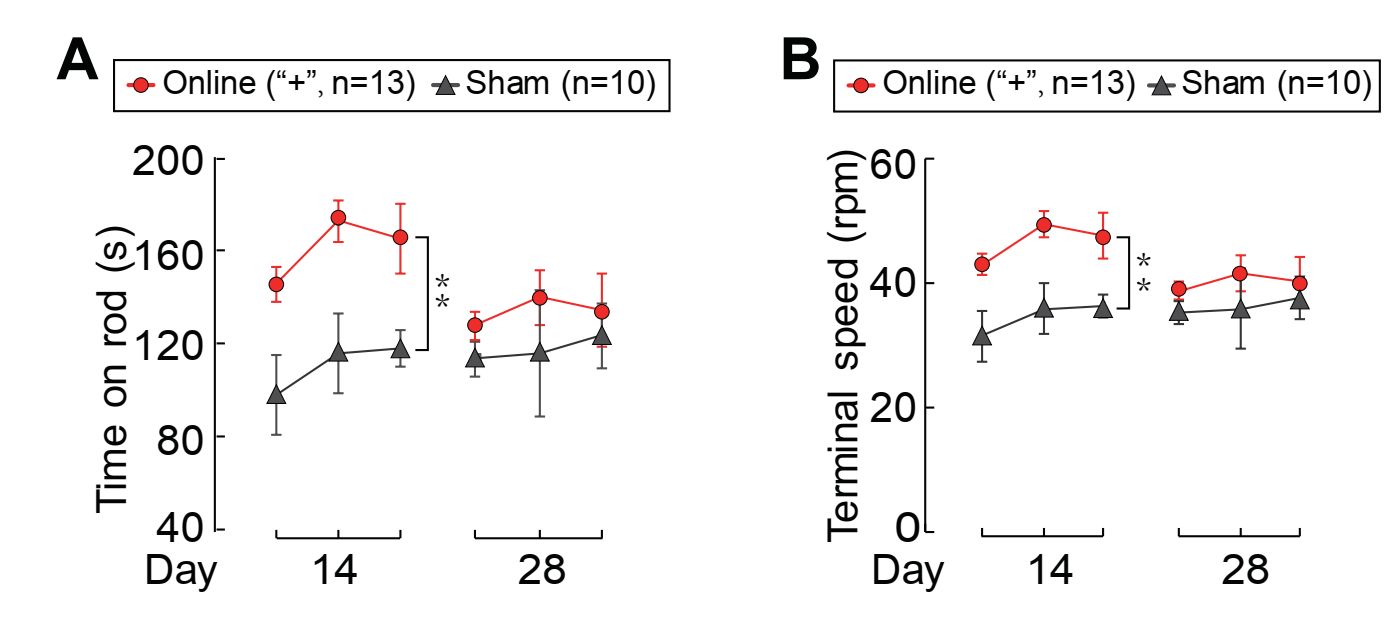

**Fig. S1 Persistence of anodal tDCS-enhanced rotarod motor learning**

(**A**) The average time of staying on the rotarod during each of the three trials, tested at 14 and 28 d after 4 d of training with the online tDCS treatment, for the same group of mice described in Fig. 1 C and D. Only the high rotation speed of 8 to 80 rpm was used in these re-tests.

(**B**) The terminal rotation speed when mice fell off the rotarod during each trial. Error bars, SEM. Significant difference was found between the data sets connected by lines (“**”, p< 0.01；two-way ANOVA).

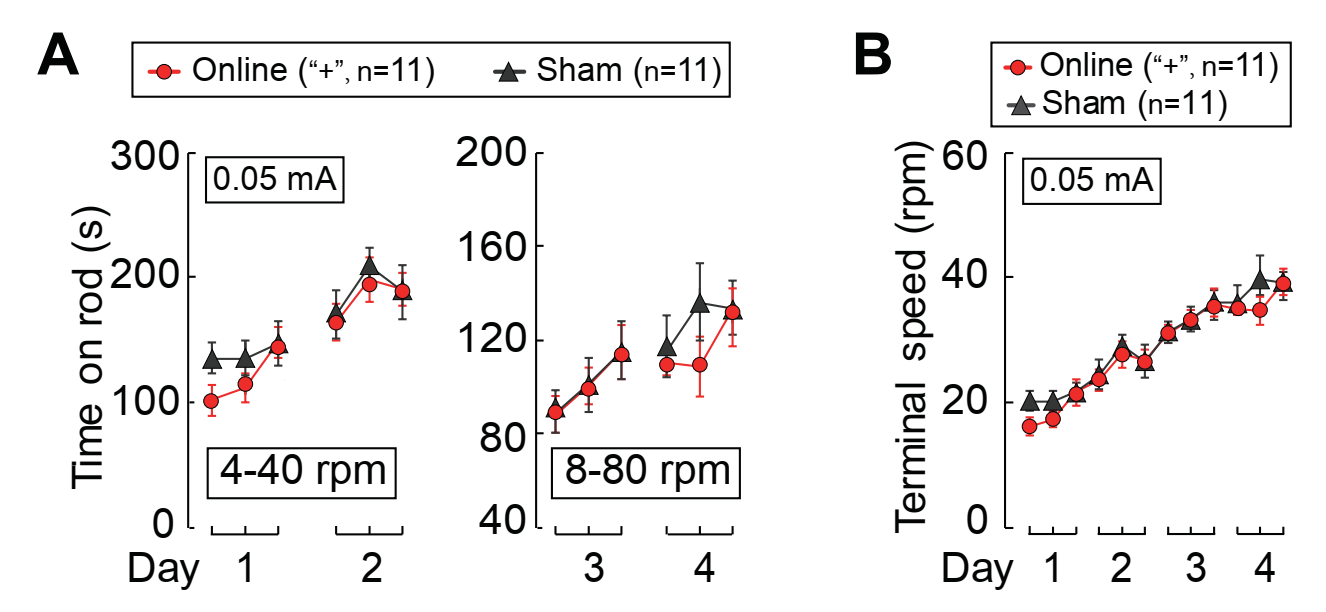

**Fig. S2** **Lower current density of anodal tDCS on primary motor cortex (M1) had no effect on motor learning**

(**A**) The average time of staying on the rotarod during each trial when mice subjected to online anodal tDCS (at 0.05 mA), using the same paradigm as that described in Fig. 1.

(**B**) The terminal rotation speed when the mouse fell from the rotarod for the same group of mice as in A. Error bars, SEM. No significant difference was found between the data sets connected by lines (two-way ANOVA).

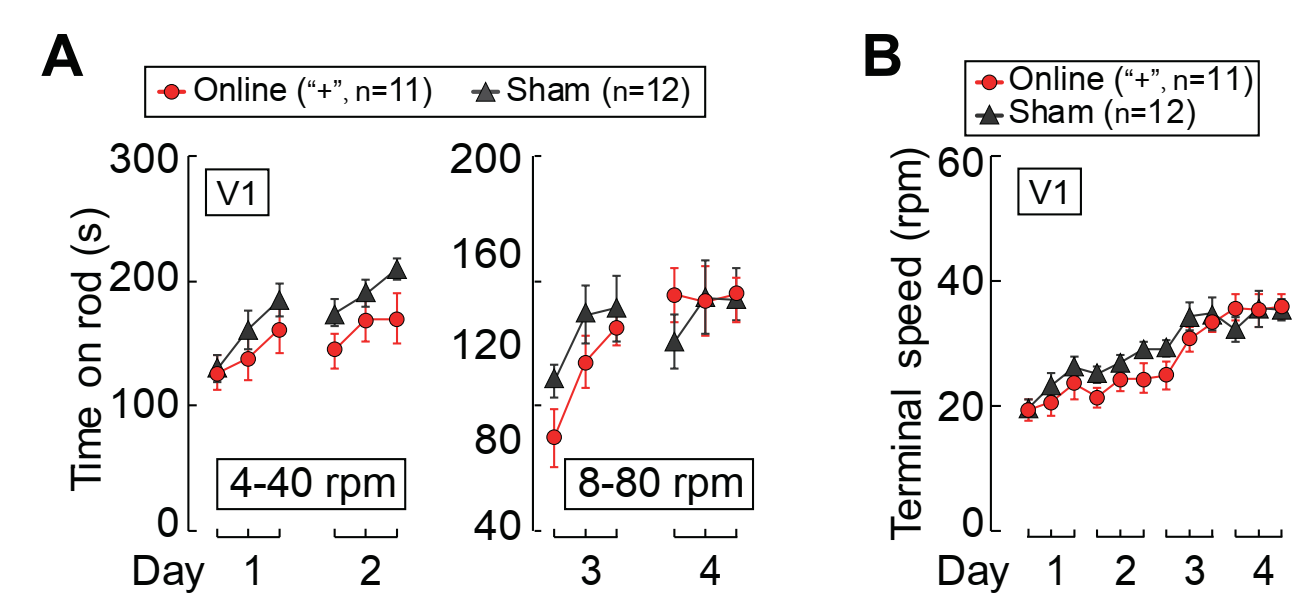

**Fig. S3 Effects of online anodal tDCS on the primary visual cortex (V1) during rotarod training**

(**A**) The average time of staying on the rotarod during each trial, with the online anodal tDCS applied to V1, using the same paradigm as that described in Fig. 1.

(**B**) The terminal rotation speed when the mouse fell from the rotarod for the same group of mice as in A. Error bars, SEM. No significant difference was found between the data sets connected by lines (two-way ANOVA).

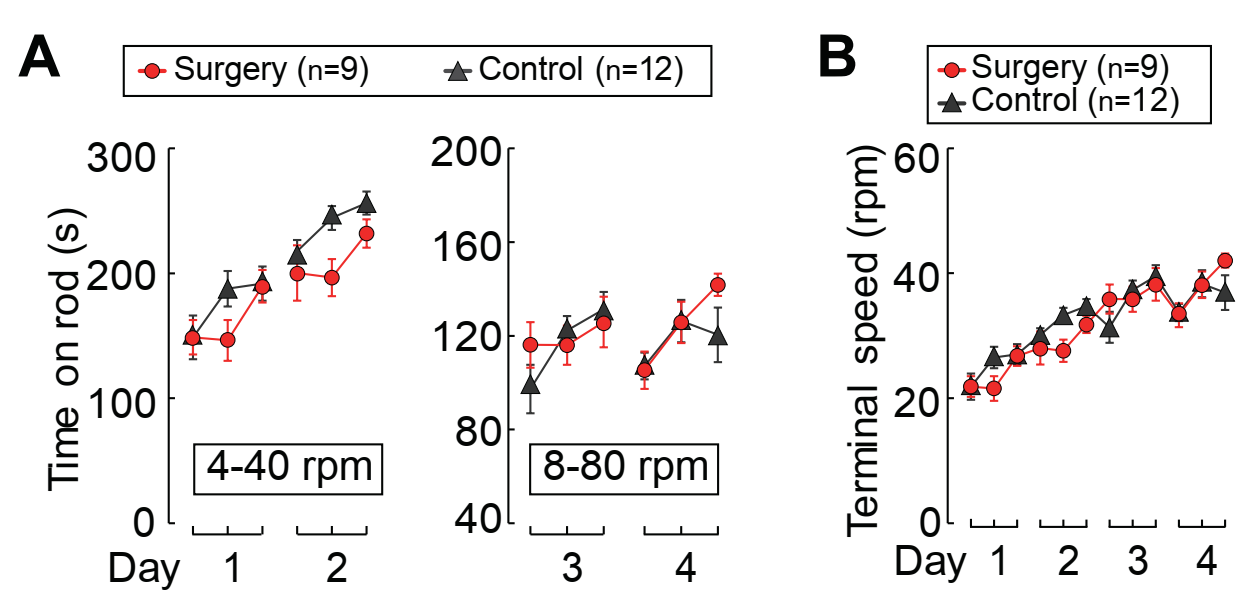

**Fig. S4 The surgery procedure of tDCS electrode implantation had no effect on mice learning the rotarod task**

(**A**) The average time of staying on the rotarod during each trial. “**Surgery**”: The mice with the surgery of tDCS electrodes implantation; “**Control**”: the mice with no surgery of tDCS electrodes implantation.

(**B**) The terminal rotation speed when mice fell from the rotarod for the same group of mice as in A. Error bars, SEM. No significant difference was found between the data sets connected by lines (two-way ANOVA).

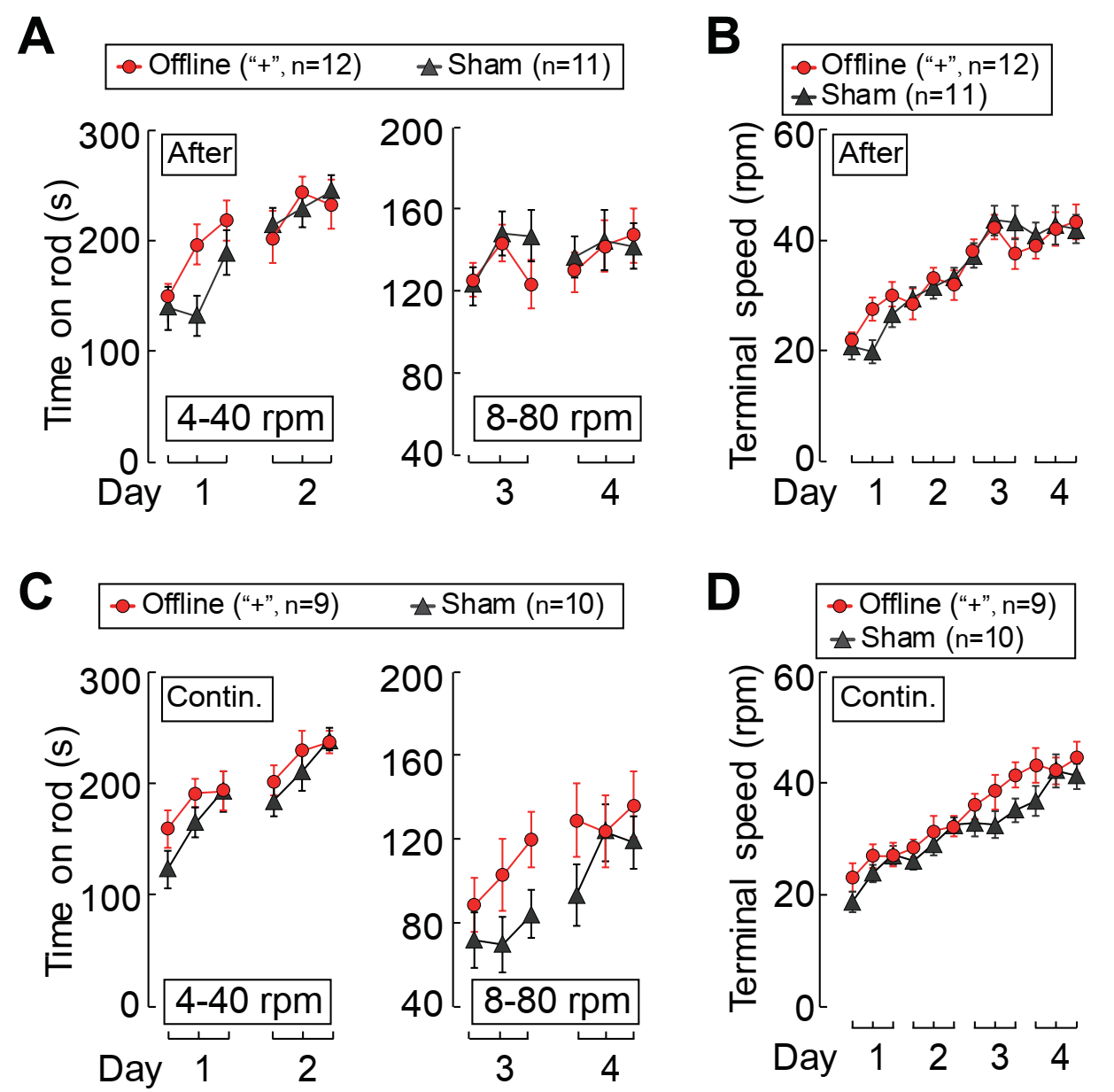

**Fig. S5 Effects of offline anodal tDCS on rotarod learning**

(**A**) The average time of staying on the rotarod during each trial when mice subjected to offline anodal tDCS (0.1 mA). “**After**”: the average values obtained with tDCS applied during ITIs after each trial.

(**B**) The terminal rotation speed when the mouse fell from the rotarod for the same group of mice as in A.

(**C**) The average time of staying on the rotarod during each trial when mice subjected to offline anodal tDCS. “**Contin.**”: 20-min tDCS continuously applied before the training.

(**D**) The terminal rotation speed when the mouse fell from the rotarod for the same group of mice as in A. Error bars, SEM. No significant difference was found between the data sets connected by lines (two-way ANOVA).

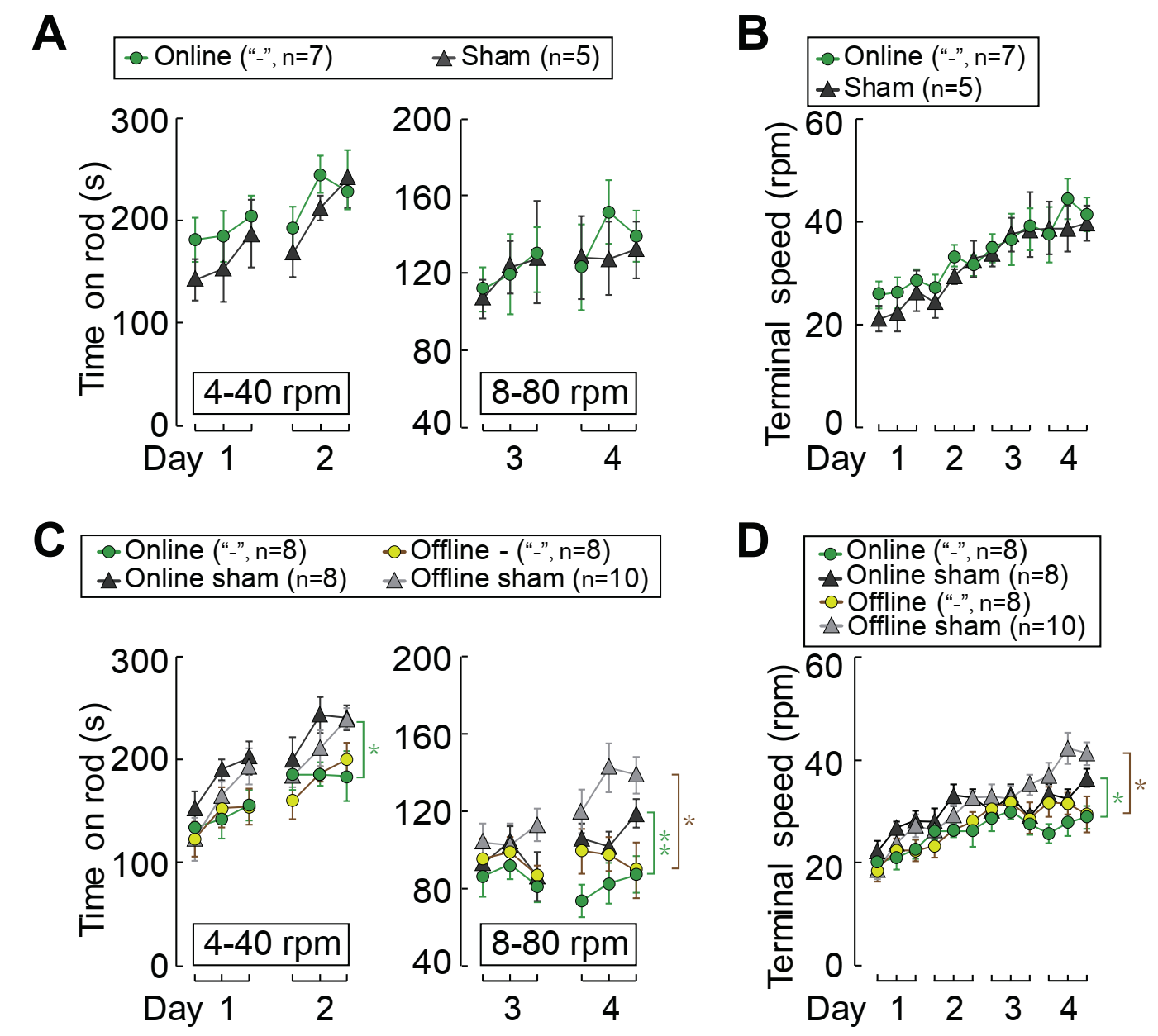

**Fig. S6 Effects of online and offline cathodal tDCS on rotarod learning** (**A**) The average time of staying on the rotarod during each trial when mice subjected to online anodal tDCS (at 0.1 mA), using the same paradigm as that described in Fig. 1.

(**B**) The terminal rotation speed when the mouse fell from the rotarod for the same group of mice as in A.

(**C**) The average time of staying on the rotarod during each trial when mice subjected to online or pre-offline cathodal tDCS (at 0.2 mA).

(**D**) The terminal rotation speed when the mouse fell from the rotarod for the same group of mice as in C. Error bars, SEM. Significant difference was found between the data sets connected by lines (“*”, p< 0.05；“**”, p< 0.01；two-way ANOVA).

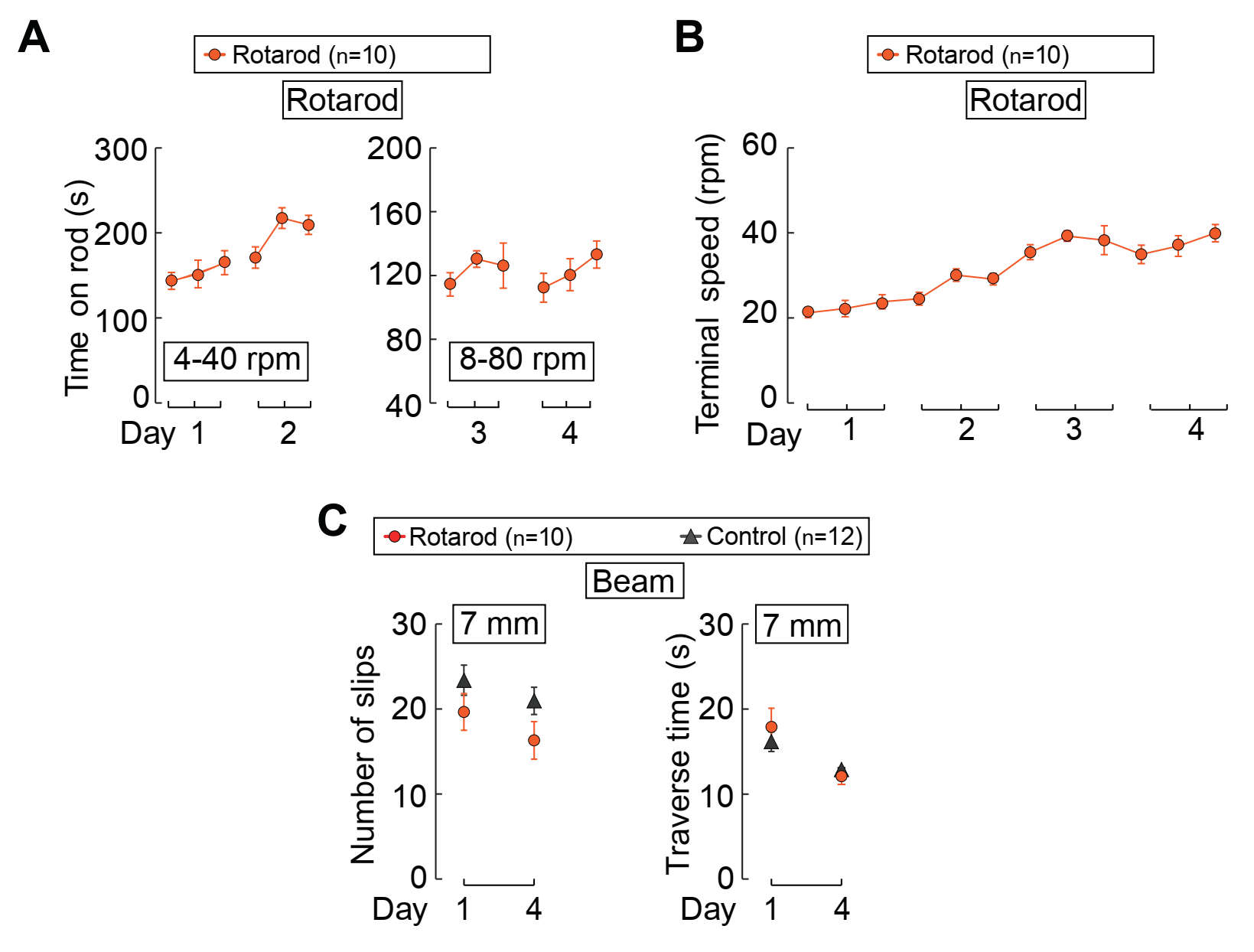

**Fig. S7 No transfer of motor learning from rotarod running to beam walking, in the absence of tDCS**

(**A**) The average time of staying on the rotarod during each trial. Mice were subjected to one trial of beam walking on a 7-mm beam before and after 4 d of rotarod training.

(**B**) The terminal rotation speed when the mice fell from the rotarod in each trial, for the same group of mice as in A.

(**C**) Data on beam walking for the “**Rotarod**” group that underwent rotarod training as described in A and B, and the “**Control**” group that were not trained for rotarod running. Number of slips and traverse time when mice walking on the beam at day 1 and day 4. Error bars, SEM. No significant difference was found between rotarod-trained group and control group ( unpaired *t* test).

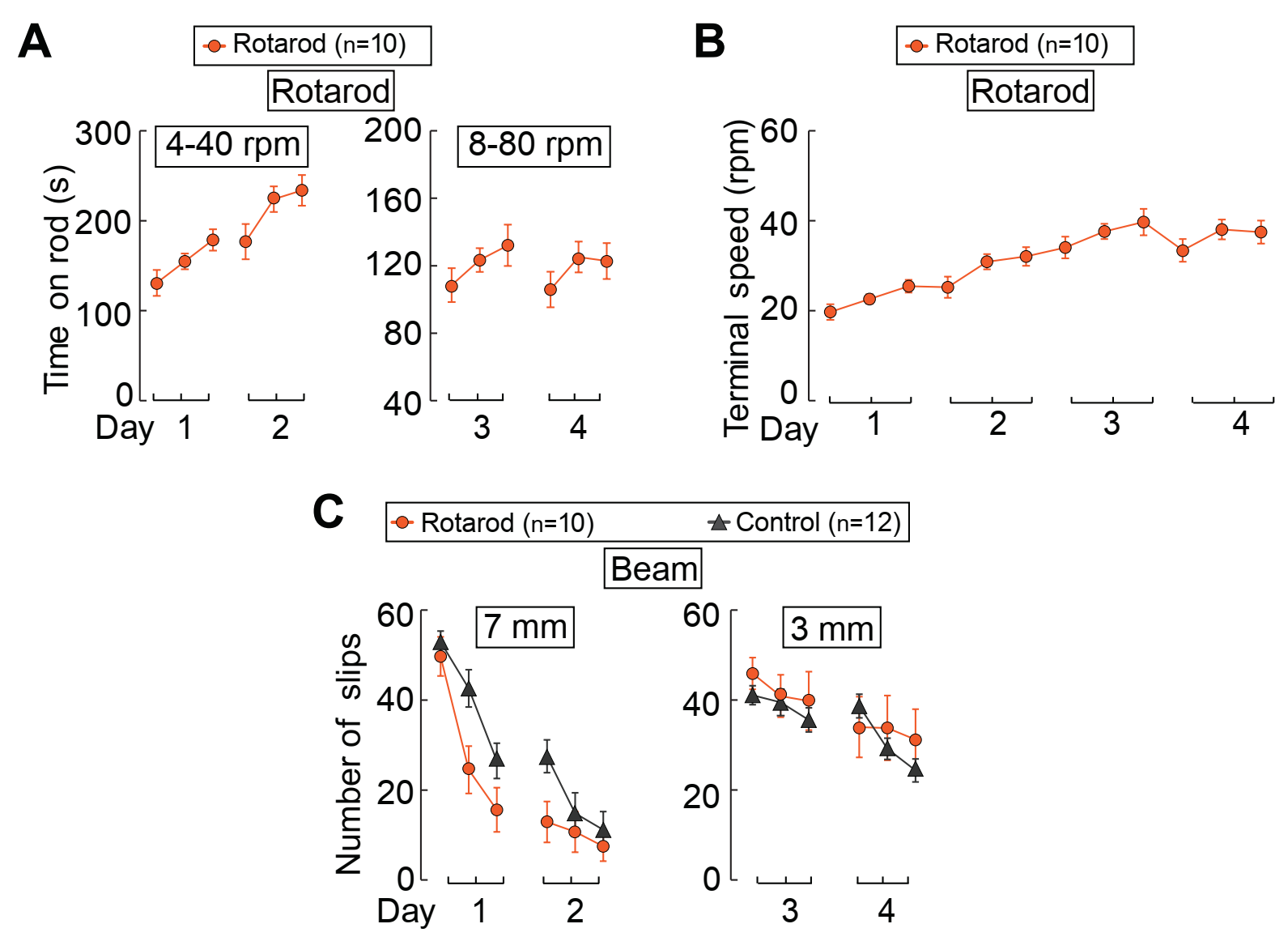

**Fig. S8 No transfer of rotarod learning to beam walking learning, in the absence of tDCS**

(**A**) The average time of staying on the rotarod during each trial. Mice were subjected to beam walking learning each day after 4-day rotarod learning.

(**B**) The terminal rotation speed when the mice fell from the rotarod in each trial, for the same group of mice as in A.

(**C**) Travers time and average number of hindlimb slips on beam walking learning for the “**Rotarod**” group that underwent rotarod training as described in A and B, and the “**Control**” group that were not trained for rotarod running. Error bars, SEM. No significant difference in beam walking learning was found between mice with and without rotarod training (two-way ANOVA).

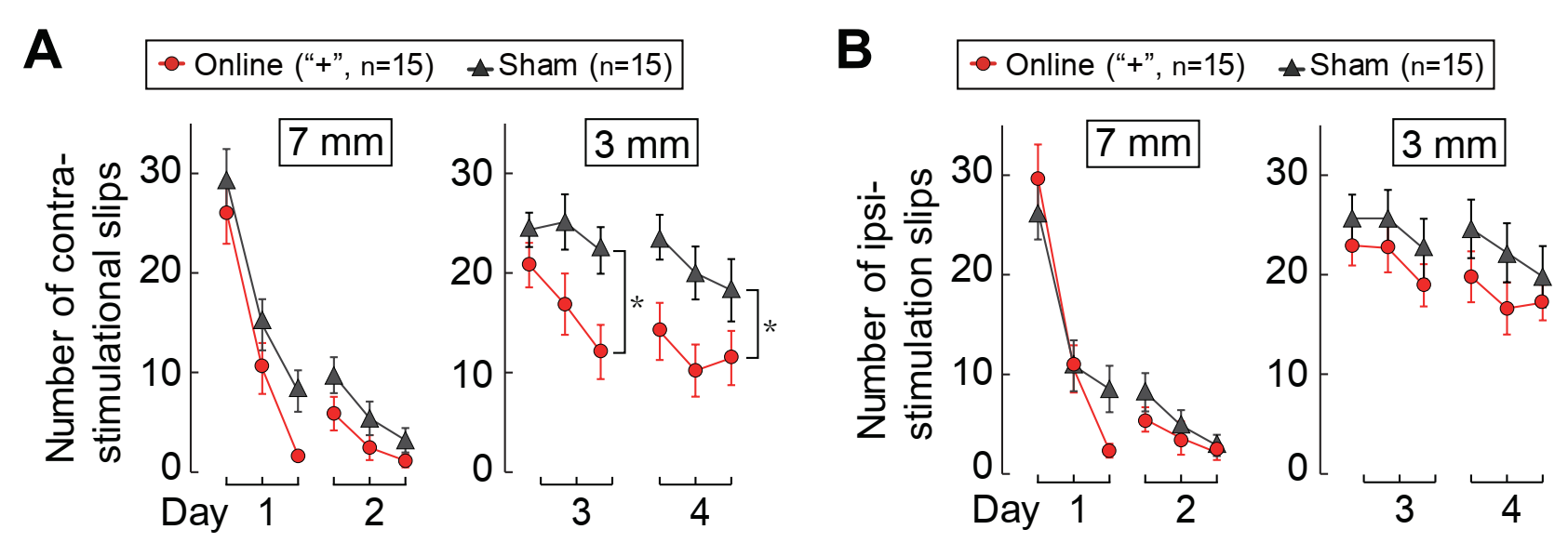

**Fig. S9 Effects of contra- and ipsi-lateral tDCS on mouse learning of beam walking task**

(**A**) The average number of contralateral hindlimb slips when mice were subjected to online anodal tDCS (at 0.1 mA) at M1.

(**B**) The average number of ipsilateral hindlimb slips when mice were subjected to the same online anodal tDCS as in A for the same group of mice. Comparing A and B, we note that online M1 stimulation reduced the number of slips only for the contralateral hindlimb. Error bars, SEM. Significant difference was found between the data sets connected by lines (“*”, p< 0.05；two-way ANOVA).

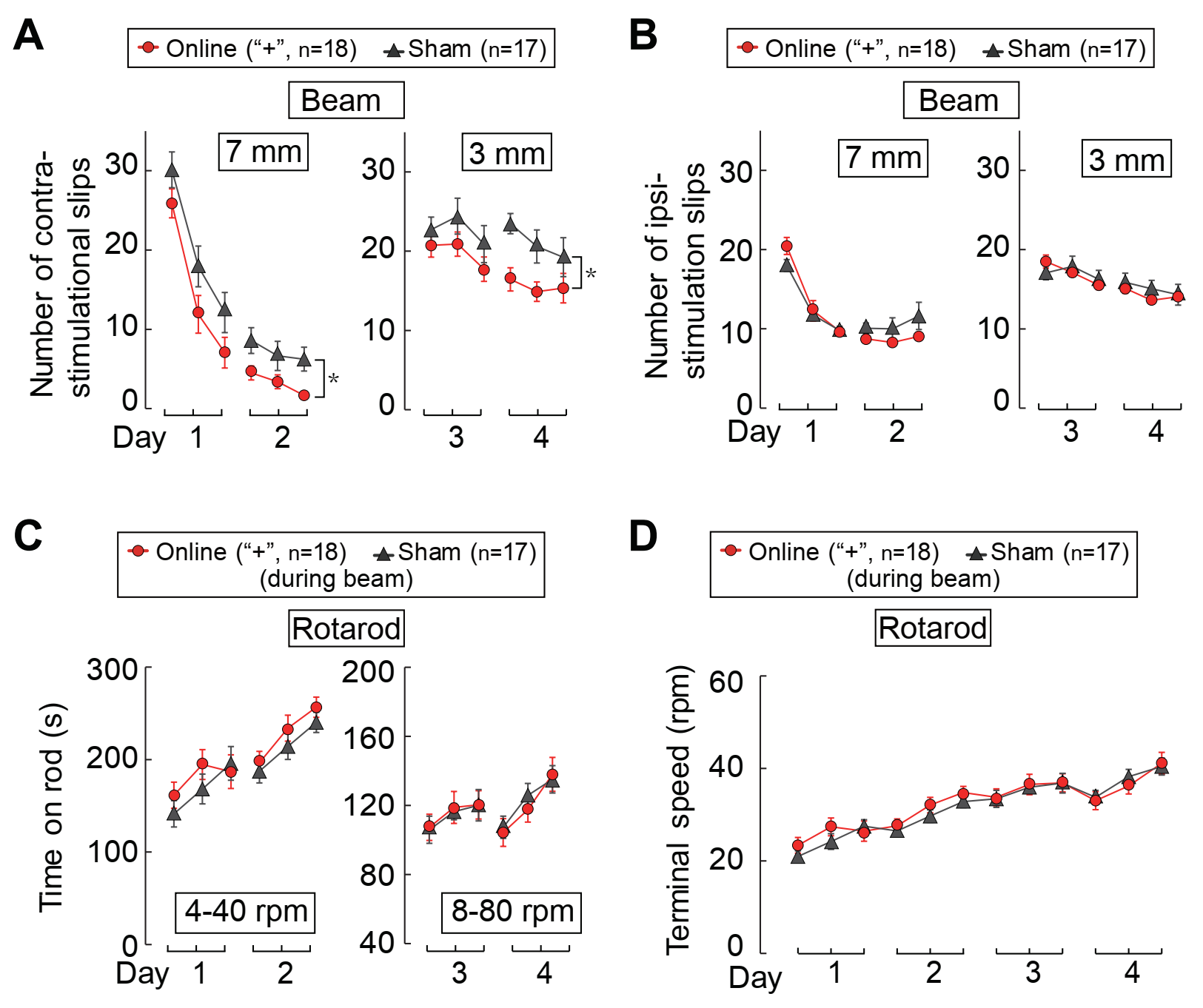

**Fig. S10 Enhancement of learning for beam walking by online anodal tDCS had no effect on rotarod learning**

(**A**) The average number of contralateral hindlimb slips when mice were subjected to online anodal tDCS (at 0.1 mA) at M1.

(**B**) The average number of ipsilateral hindlimb slips when mice were subjected to online anodal tDCS for the same group of mice as in A.

(**C**) The beam walking learning was followed by a rotarod training task each day in the absence of tDCS. The average time of staying on the rotarod during each trial followed by beam walking learning for the same group mice as in A.

(**D**) The terminal speed when mice fell from the rotarod for the same group mice as in A. Error bars, SEM. Significant difference was found between the data sets connected by lines (“*”, p< 0.05；two-way ANOVA).

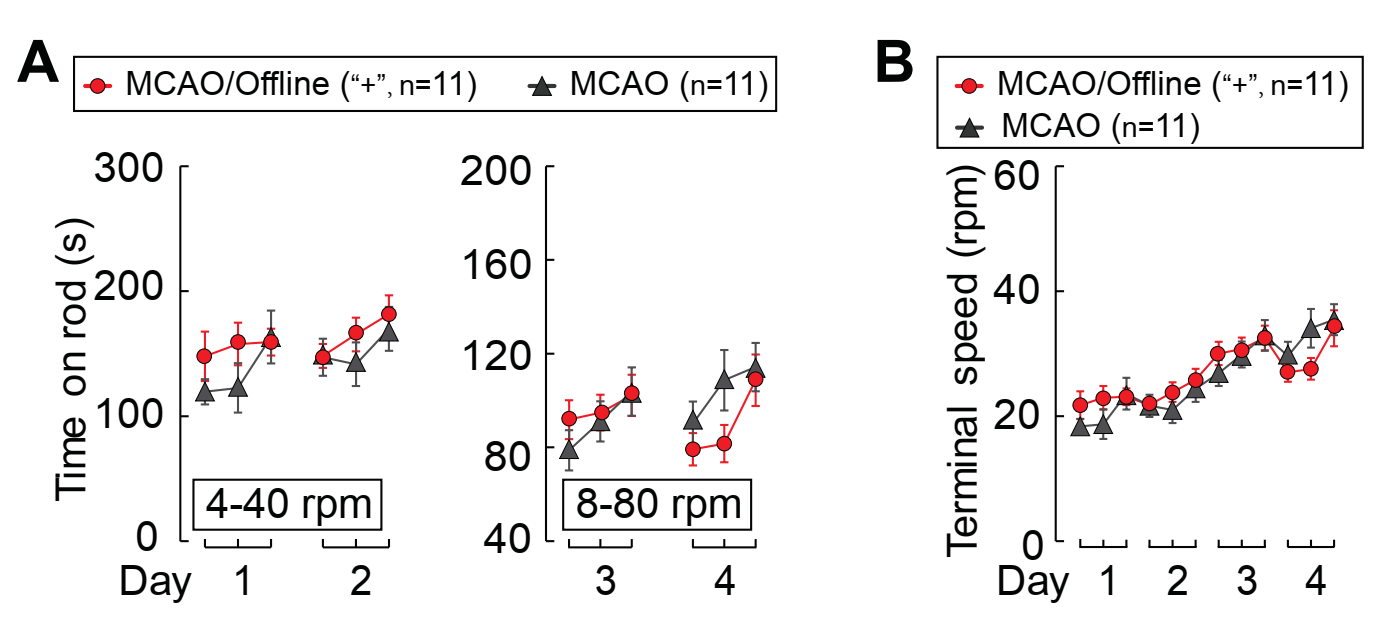

**Fig. S11 No effects of offline anodal tDCS on rotarod learning of MCAO mice**

(**A**) The average time of staying on the rotarod during each trial when MCAO stroke model mice were subjected to offline anodal tDCS at M1 ipsilateral to the infarct site.

(**B**) The terminal rotation speed when the mouse fell from the rotarod for the same group of mice as in A. Error bars, SEM. No significant difference was found between the data sets connected by lines (two-way ANOVA).

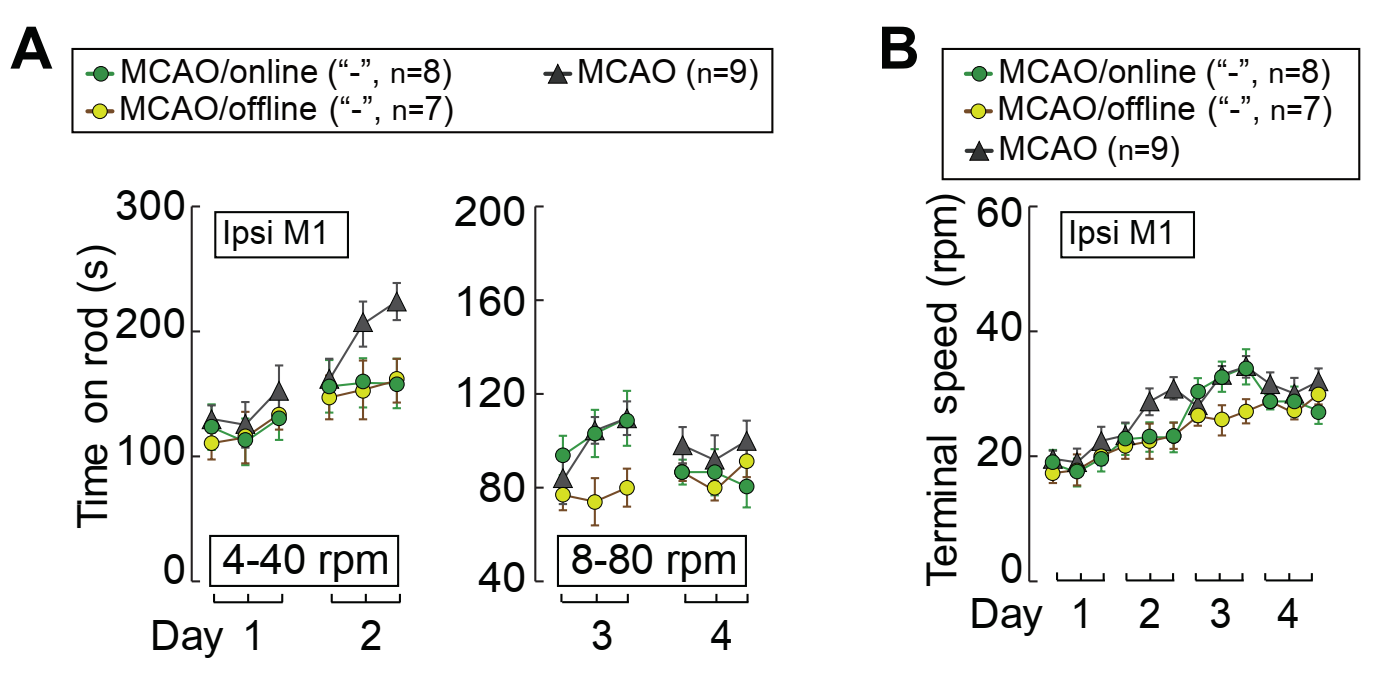

**Fig. S12 Effects of online or offline cathodal tDCS on rotarod learning of MCAO mice**

(**A** and **B**) The average time of staying on the rotarod (A) and the terminal speed when the MCAO mouse fell from the rotarod (B) during each trial when MCAO stroke model mice subjected to online or offline cathodal tDCS (at 0.2 mA) of M1 ipsilateral to the infarct site (“**Ipsi M1**”). Error bars, SEM. No significant difference was found between the data sets connected by lines (two-way ANOVA).

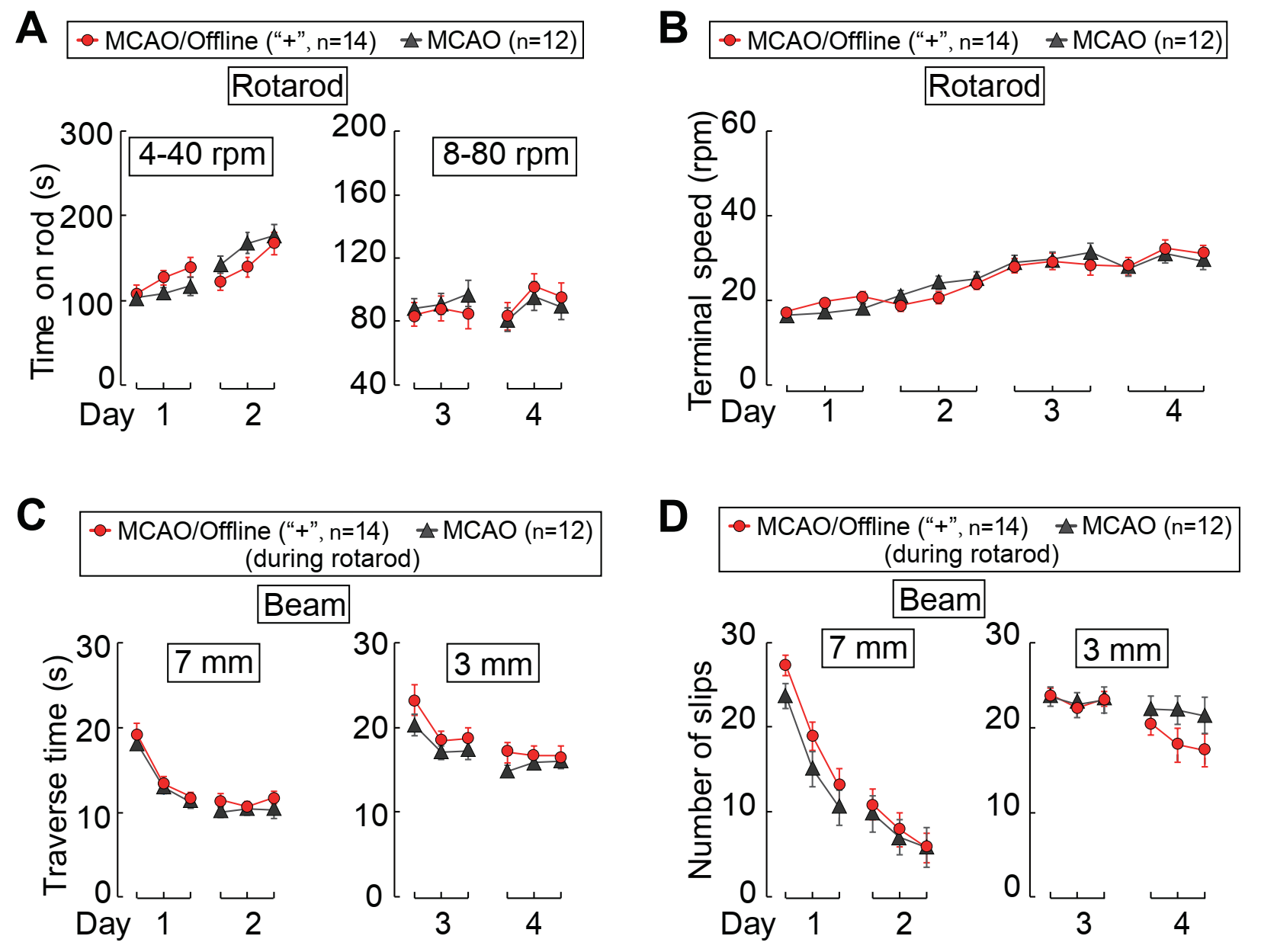

**Fig. S13 No effect of offline anodal tDCS on either rotarod or beam walking learning for MCAO stroke mice performing dual-task training**

(**A** and **B**) Data depicting rotarod learning, with offline anodal tDCS at M1 ipsilateral to the infarct site, using the same dual-task paradigm as described in Fig. 2.

(**C** and **D**) Data depicting beam walking learning, as measured by the traverse time and contralateral hindlimb slips, for the same group of mice as in A. Error bars, SEM. No significant difference was found between the data sets connected by lines (two-way ANOVA).

**Movies Attachment**

**Movies S1 and S2.** Video images of mice performing rotarod running at day 1 (S1) and day 4 (S2) of training. Each movie shows one mouse subjected to online anodal tDCS (0.1 mA), and the other subjected to sham treatment (0 mA).

**Movies S3 to S6.** Video images of mice performing beam walking at day 1 (S3, S5) and day 4 (S4, S6) of training. The movies show the performance of the same mouse that was subjected to the online anodal tDCS (0.1 mA, S3 and S4) and another mouse that was subjected to the sham treatment (0 mA, S5 and S6).

**Movies S7 and S8.** Video images of fluorescence changes in a population of M1 neurons in a Thy-1 transgenic mouse expressing GCaMP6s in response to two episodes of anodal (S7) and cathodal (S8) tDCS, as monitored by *in vivo* two-photon imaging of a head-fixed mouse running on the rotating treadmill.

**Movies S9 and S10.** Video images of MCAO stroke mice performing rotarod running at day 1 (S1) and day 4 (S2) of training. Each movie shows one mouse subjected to online anodal tDCS (0.1 mA), and the other subjected to sham treatment (0 mA).

PASS 15.0.5

**Two-Sample T-Tests Allowing Unequal Variance**

**Numeric Results for Two-Sample T-Test Allowing Unequal Variance**

Alternative Hypothesis: H1: δ = μ1 - μ2 ≠ 0

| **Target**  **Power** | **Actual**  **Power** | **N1** | **N2** | **N** | **μ1** | **μ2** | **δ** | **σ1** | **σ2** | **Alpha** |
| --- | --- | --- | --- | --- | --- | --- | --- | --- | --- | --- |
| 0.9 | 1.00000 |  |  | 0 | 0.5 | 0.5 | 0.0 | 0.3 | 0.3 | 0.050 |
| 0.9 | 0.91255 | 9 | 9 | 18 | 0.5 | 1.0 | -0.5 | 0.3 | 0.3 | 0.05 |
| 0.9 | 0.97266 | 4 | 4 | 8 | 0.5 | 1.5 | -1.0 | 0.3 | 0.3 | 0.05 |
| 0.9 | 0.99278 | 3 | 3 | 6 | 0.5 | 2.0 | -1.5 | 0.3 | 0.3 | 0.05 |
| 0.9 | 0.99992 | 3 | 3 | 6 | 0.5 | 2.5 | -2.0 | 0.3 | 0.3 | 0.05 |
| 0.9 | 0.91255 | 9 | 9 | 18 | 1.0 | 0.5 | 0.5 | 0.3 | 0.3 | 0.05 |
| 0.9 | 0.91255 |  |  | 0 | 1.0 | 1.0 | 0.0 | 0.3 | 0.3 | 0.05 |
| 0.9 | 0.91255 | 9 | 9 | 18 | 1.0 | 1.5 | -0.5 | 0.3 | 0.3 | 0.05 |
| 0.9 | 0.97266 | 4 | 4 | 8 | 1.0 | 2.0 | -1.0 | 0.3 | 0.3 | 0.05 |
| 0.9 | 0.99278 | 3 | 3 | 6 | 1.0 | 2.5 | -1.5 | 0.3 | 0.3 | 0.05 |
| 0.9 | 0.97266 | 4 | 4 | 8 | 1.5 | 0.5 | 1.0 | 0.3 | 0.3 | 0.05 |
| 0.9 | 0.91255 | 9 | 9 | 18 | 1.5 | 1.0 | 0.5 | 0.3 | 0.3 | 0.05 |
| 0.9 | 0.91255 |  |  | 0 | 1.5 | 1.5 | 0.0 | 0.3 | 0.3 | 0.05 |
| 0.9 | 0.91255 | 9 | 9 | 18 | 1.5 | 2.0 | -0.5 | 0.3 | 0.3 | 0.05 |
| 0.9 | 0.97266 | 4 | 4 | 8 | 1.5 | 2.5 | -1.0 | 0.3 | 0.3 | 0.05 |

Edition. Blackwell Science. Malden, MA.

Zar, Jerrold H. 1984. Biostatistical Analysis (Second Edition). Prentice-Hall. Englewood Cliffs, New Jersey.

**Report Definitions**

Target Power is the desired power value (or values) entered in the procedure. Power is the probability of

rejecting a false null hypothesis.

Actual Power is the power obtained in this scenario. Because N1 and N2 are discrete, this value is often

(slightly) larger than the target power.

N1 and N2 are the number of items sampled from each population.

N is the total sample size, N1 + N2.

μ1 and μ2 are the assumed population means.

δ = μ1 - μ2 is the difference between population means at which power and sample size calculations are made.

σ1 and σ2 are the assumed population standard deviations for groups 1 and 2, respectively.

Alpha is the probability of rejecting a true null hypothesis.

**Summary Statements**

Group sample sizes of NA and NA achieve 100.000% power to reject the null hypothesis of equal

means when the population mean difference is μ1 - μ2 = 0.5 - 0.5 = 0.0 with standard deviations

of 0.3 for group 1 and 0.3 for group 2, and with a significance level (alpha) of 0.050 using a

two-sided two-sample unequal-variance t-test.

PASS 15.0.5

**Two-Sample T-Tests Allowing Unequal Variance**

**Dropout-Inflated Sample Size**

**Dropout-Inflated	Expected**

**Enrollment	Number of**

**──── Sample Size ────	──── Sample Size ────	───── Dropouts ─────**

**Dropout Rate	N1	N2	N	N1'	N2'	N'	D1	D2	D**

20%	0

20%	9	9	18	12	12	24	3	3	6

20%	4	4	8	5	5	10	1	1	2

20%	3	3	6	4	4	8	1	1	2

20%	3	3	6	4	4	8	1	1	2

20%	9	9	18	12	12	24	3	3	6

20%	0

20%	9	9	18	12	12	24	3	3	6

20%	4	4	8	5	5	10	1	1	2

20%	3	3	6	4	4	8	1	1	2

20%	4	4	8	5	5	10	1	1	2

20%	9	9	18	12	12	24	3	3	6

20%	0

20%	9	9	18	12	12	24	3	3	6

20%	4	4	8	5	5	10	1	1	2

**Definitions**

Dropout Rate (DR) is the percentage of subjects (or items) that are expected to be lost at random during the

course of the study and for whom no response data will be collected (i.e. will be treated as "missing").

N1, N2, and N are the evaluable sample sizes at which power is computed. If N1 and N2 subjects are evaluated

out of the N1' and N2' subjects that are enrolled in the study, the design will achieve the stated power.

N1', N2', and N' are the number of subjects that should be enrolled in the study in order to end up with N1,

N2, and N evaluable subjects, based on the assumed dropout rate. After solving for N1 and N2, N1' and N2'

are calculated by inflating N1 and N2 using the formulas N1' = N1 / (1 - DR) and N2' = N2 / (1 - DR), with

N1' and N2' always rounded up. (See Julious, S.A. (2010) pages 52-53, or Chow, S.C., Shao, J., and Wang, H.

(2008) pages 39-40.)

D1, D2, and D are the expected number of dropouts. D1 = N1' - N1, D2 = N2' - N2, and D = D1 + D2.

PASS 15.0.5

**Two-Sample T-Tests Allowing Unequal Variance**

**Chart Section**

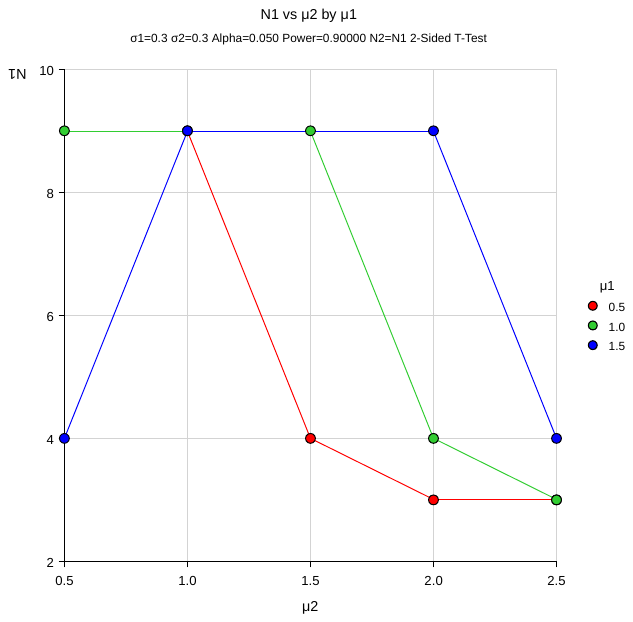

PASS 15.0.5

**Two-Sample T-Tests Allowing Unequal Variance**

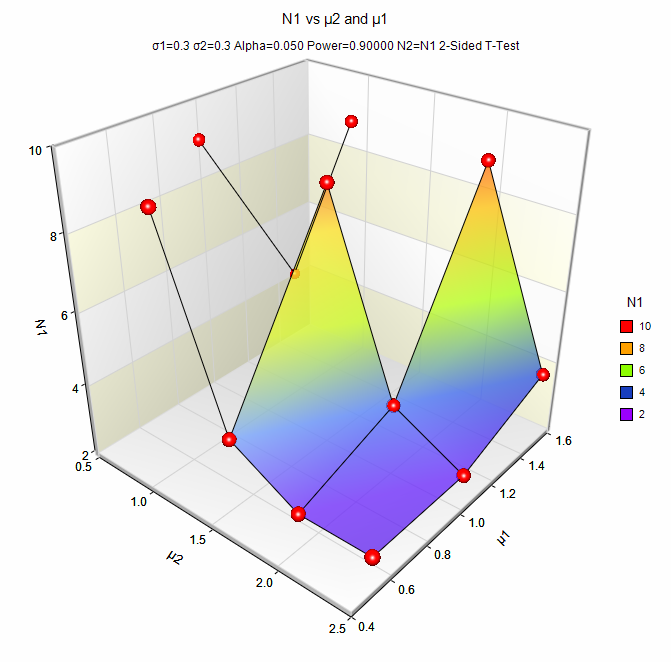

PASS 15.0.5

**Two-Sample T-Tests Allowing Unequal Variance**

**Procedure Input Settings**

**Autosaved Template File**

C:\Users\pc\Documents\PASS 15\Procedure Templates\Autosave\Two-Sample T-Tests Allowing Unequal Variance - Autosaved 2022_3_16-15_17_16.t389

**Design Tab**

Solve For:	Sample Size

Alternative Hypothesis:	Two-Sided

Power:	0.9

Alpha:	0.05

Group Allocation:	Equal (N1 = N2)

Input Type:	Means

μ1:	0.5 to 1.5 by 0.5

μ2:	0.5 to 2.5 by 0.5

σ1:	0.3

σ2:	0.3

The power for predicting the sample size of experimental results has been calculated for statistical analysis. According to previous relevant studies and our study about rotarod, the standard deviation (σ) is usually under 0.3, the mean of control group is usually between 0.5 to 1.5 and the mean of experimental group is usually between 0.5 to 2.5. The power is designed at 90%, which is considered to be largely valid. From the results of power calculation (by PASS, NCSS Inc.), our experiments satisfy the minimum sample size.
